# Split-gene drive system provides flexible application for safe laboratory investigation and potential field deployment

**DOI:** 10.1101/684597

**Authors:** Víctor López Del Amo, Alena L. Bishop, Héctor M. Sánchez C., Jared B. Bennett, Xuechun Feng, John M. Marshall, Ethan Bier, Valentino M. Gantz

## Abstract

CRISPR-based gene drives spread through populations bypassing the dictates of Mendelian genetics, offering a population-engineering tool for tackling vector-borne diseases, managing crop pests, and helping island conservation efforts; unfortunately, current technologies raise safety concerns for unintended gene propagation. Herein, we address this by splitting the two drive components, Cas9 and gRNAs, into separate alleles to form a novel trans-complementing split–gene-drive (tGD) and demonstrate its ability to promote super-Mendelian inheritance of the separate transgenes. This bi-component nature allows for individual transgene optimization and increases safety by restricting escape concerns to experimentation windows. We employ the tGD and a small– molecule-controlled version to investigate the biology of component inheritance and use our system to study the maternal effects on CRISPR inheritance, impaired homology on efficiency, and resistant allele formation. Lastly, mathematical modeling of tGD spread in a population shows potential advantages for improving current gene-drive technologies for field population modification.

## INTRODUCTION

CRISPR gene-drive systems have a tremendous potential for wild population engineering due to their ability to self-propagate, biasing inheritance from Mendelian (50%) to super-Mendelian (>50%)^1–7^. This technology is promising for fighting vector-borne diseases (e.g. malaria) by suppressing^3,4^ or modifying mosquito populations^5^ to decrease their burden on public health. Additionally, the technology can be used to manage crop pests^8,9^ and suppress invasive rodents in island restoration efforts^10,11^. While the scientific community welcomes this technology’s enormous potential for solving significant global issues, it also acknowledges that it is currently in its infancy and that several gaps need to be filled before it can be safely deployed^12–14^. In particular, concerns have been raised about accidental release during laboratory research, begging for the development of strategies to increase safety during optimization phases^15^.

Gene-drive systems use an allelic conversion process that occurs in the germline, changing heterozygous to homozygous cells that can achieve the super-Mendelian inheritance necessary for population engineering. Currently, two different approaches based on the RNA-guided endonuclease Cas9 are feasible: **1)** a **full gene drive (full-GD)** is the traditional format, consisting of a Cas9 and a guide RNA (gRNA) gene inserted at the target location as a single unit. The two gene products combine to induce a double strand break at the same position on the wild-type allele, which is then repaired via homology directed repair (HDR) using the intact chromosome carrying the gene-drive element as a template^2–5,16–18^. **2)** The **gRNA-only gene drive (gRNA-GD)** is based on CopyCat gRNA elements^19^ that are capable of allelic conversion in the presence of a genetic source of Cas9. Since only the gRNA element is propagated in this case, its spread is regulated by the presence of separate, static Cas9 transgene^10,20,21^. The use of a full-GD is causing concern to the scientific community as an accidental release could spread unchecked^15^. While a gRNA-GD would address such concerns, its application in the field for large scale population engineering is unlikely since it would require a large percentage of the population carrying a Cas9 transgene^22^.

Here, we develop a novel CRISPR gene-drive method in *Drosophila* called **trans-complementing gene drive (tGD)**, which combines the strengths of both approaches described above. This arrangement splits the Cas9 and gRNA into two different transgenic lines that, when combined by genetic cross, exhibit the same properties of a full-GD, as both elements are capable of propagating. The scope of this work is to demonstrate that tGD can bias the inheritance of two interdependent transgenes and that it can be used to study the impact of specific transgene parameters, such as the here-described effects of Cas9 promoter, maternal effect, genomic context, and homology, on the gene drive efficiency. Additionally, we apply a drug-regulation technology to the tGD system such that super-Mendelian inheritance can be controlled by the presence of a small molecule in the fly diet, and use this to study Cas9 activation in the adult germline. Lastly, we simulate the propagation of tGD elements and uncover their potential to spread transgenes to a higher fraction of a population than a corresponding full-drive system, highlighting the tGD’s potential for future field applications.

## RESULTS AND DISCUSSION

### The tGD system displays super-Mendelian behavior

The tGD system was designed to split the two genetic elements, Cas9 and a two-part gRNA gene construct (gRNA-A and gRNA-B), into two distinct genomic locations that, when separated, would behave as regular transgenes (no gene-drive activity) (**Fig. 1A**). After assembly through genetic cross, gRNA-A would cut at the Cas9 site while gRNA-B would cut at the gRNA site (**Fig. 1A**). Since the cleaved ends would have perfect homology to the sequences flanking each of the transgenic elements, the HDR pathway would insert a copy of each transgene into the wild-type allele (**Fig. 1A**).

**Figure 1.**
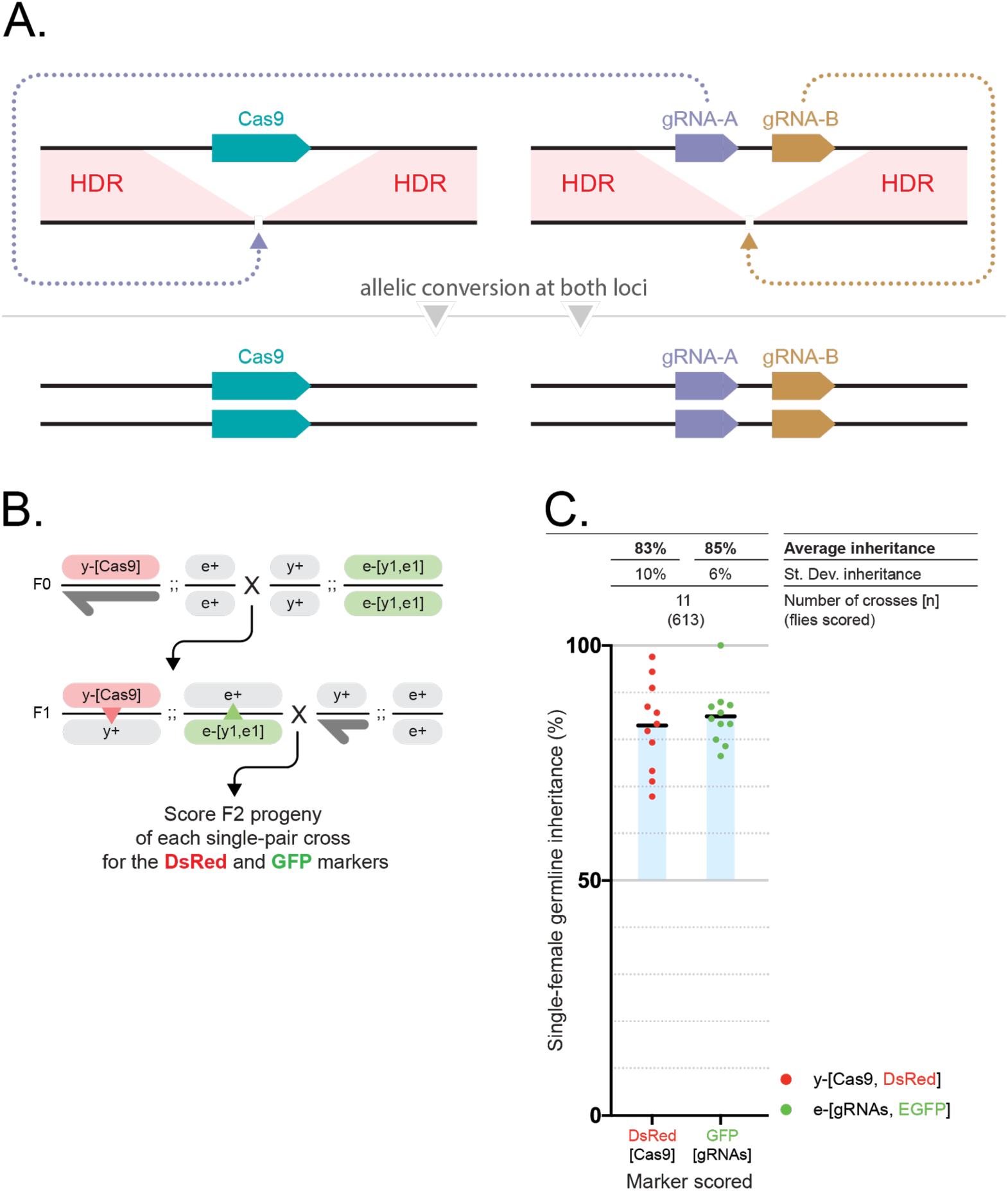
The trans-complementing gene-drive system (tGD) features simultaneous super-Mendelian inheritance of two transgenes. (**A**) Schematic of the tGD genetic arrangement with two elements that can be kept separated as different transgenic lines. The Cas9 transgene is inserted in the genomic location targeted by the gRNA-A, while a separate cassette expressing a tandem gRNA construct (gRNA-A, gRNA-B) is inserted at the location targeted by gRNA-B. Upon genetic cross, each gRNA combines with Cas9 to generate a double-strand DNA break at each locus on the wild-type allele. Each break is then repaired by homology-directed repair (HDR) pathway using the intact chromosome carrying the transgene as a template. (**B**) Outline of the genetic cross used to demonstrate tGD in fruit flies. F0 males carrying a DsRed-marked Cas9 transgene inserted in the *yellow* locus were crossed to females carrying a GFP-marked cassette containing two gRNAs (*y*1-*e*1) inserted in the *ebony* coding sequence. Trans-heterozygous F1 females (carrying both Cas9 and gRNAs) were crossed to wild-type males to assess germline transmission rates of the fluorophores marking the transgenes in the F2 progeny. The conversion event is indicated by the red and green triangles in the F1 females (**C**) Single F1 female germline inheritance output measured as GFP and DsRed marker presence in the F2 progeny. The black bar represents the inheritance average. The blue shading represents the deviation from the expected 50% “Mendelian” inheritance. Inheritance average, standard deviation, number of samples (n) and total number of flies scored in each experiment are represented over the graph in line with the respective data.

To form this system, we first generated a tGD targeting the *yellow* (*y*) and *ebony* (*e*) coding sequences (tGD(y,e)), as both genes have a whole-body display of their lighter and darker body color phenotype, respectively, and are therefore easily detectable^23^. To do this, we introduced a *Streptococcus pyogenes Sp*Cas9 (Cas9) source driven by the *vasa* promoter into the *yellow* gene (X chromosome). We marked Cas9 with a DsRed (Red) fluorescent reporter expressed in the eye to generate the *vasa*-Cas9 line (**Supplementary Fig. 1**). We placed the second transgene, carrying the gRNA tandem cassette (gRNA-*y1* and gRNA-*e1*), on chromosome III (autosome), disrupting the *ebony* gene. This cassette was instead marked with EGFP (Green) to generate the *e*-[*y1,e1*] line (**Supplementary Fig. 1**).

To test this tGD(*y,e*) arrangement, we individually crossed *vasa-*Cas9 males to *e*-[*y1,e1*] virgin females (F0, **Fig. 1B**), collected F1 trans-heterozygous virgin females carrying both constructs, and single-pair crossed them to individual males of our wild-type genetic background strain, Oregon-R (Or-R) (F1; **Fig. 1B**). The phenotypical analysis of the fluorescent markers in the resulting F2 progeny allowed simultaneous evaluations of the germline output inheritance rates of both the Cas9 and gRNA transgenes of each single F1 female (F2; **Fig. 1B**). We scored the F2 progeny of 11 F1 females and observed an inheritance greater than 50% for either transgene, with an average of 83% and 85% for Cas9-Red and gRNA-Green, respectively (**Fig. 1C, Supplementary Data 1**). These results demonstrate for the first time that a CRISPR gene drive can be split into two separate genetic elements located on different chromosomes and that they can be simultaneously propagated with super-Mendelian inheritance, offering flexibility and safety while functioning as a full-GD.

We next confirmed this tGD functionality using a separate, previously-validated locus, *white* (*w*)^1,2,17^. To test the tGD(y,w), we generated an EGFP-tagged tandem-gRNA element targeting both *y* and *w* to generate a *w*-[*y1,w1*] line (**Supplementary Fig. 1**). These *w*-[*y1,w1*] virgin females (F0) were crossed with *y*-inserted *vasa*-Cas9 males, and the F1 virgin females were collected and outcrossed to Or-R males (**Fig. 2A**). In the F2 progeny, we observed 89% and 96% inheritance rates of the Cas9-Red and gRNAs-Green transgenes, respectively (**Fig. 2E**), which were both higher than when the gRNA construct was inserted in *ebony*. This suggested that genomic location might affect the allelic conversion efficiency, so we tested this by switching the location of both transgenes, placing the Cas9 element in *white* and the gRNAs in *yellow* to generate tGD(w,y) (**Supplementary Fig. 1**). This switched version displayed 95% and 98% conversion efficiency for the Cas9 and gRNA constructs (**Supplementary Fig. 2, Supplementary Data 2**), alleviating the significant difference between the two transgenes that was observed for tGD(y,w) (89% at *yellow* and 96% at *white*) and offering higher conversion rates than previous studies using a full-GD at the *y* and *w* loci^2,17^. This discrepancy could be due to the different gRNA efficiencies^24,25^ as well as the use of w[118] mutant or Canton-S strains in previous studies versus our Or-R, wherein differences in genetic background may affect gene drive efficiency^2^. Regardless, these results suggest that conversion efficiencies are impacted by the genomic location of the transgene and that transgene size in the range tested (3–8.3 kbp) does not negatively affect allelic conversion in our system.

**Figure 2.**
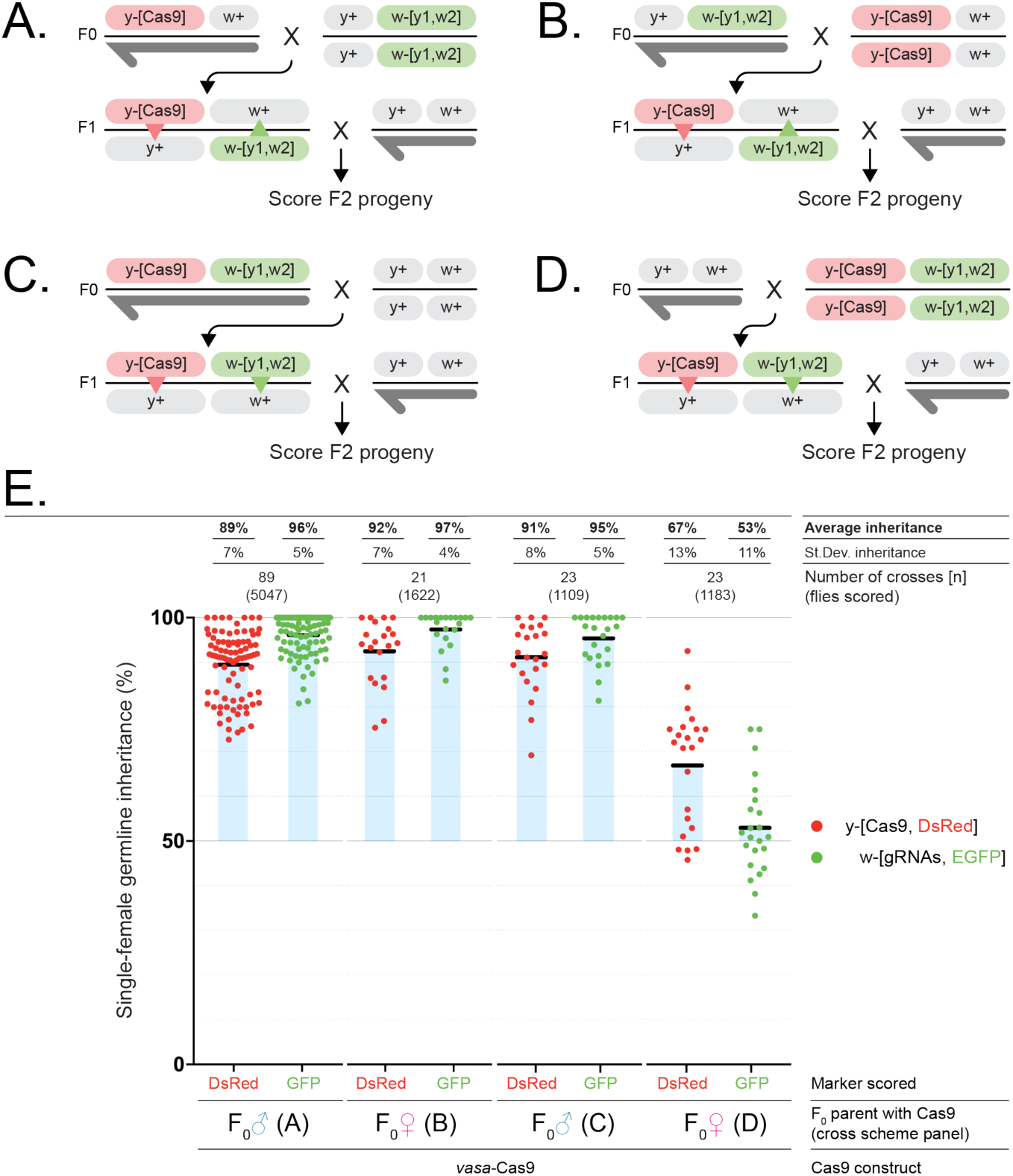
tGD targeting the *yellow* and *white* loci uncovers maternal effect mechanism. (**A-D**) Genetic crosses performed using the tGD elements targeting *yellow* (DsRed-Cas9, as Fig. 1) and *white* (GFP-y1,w2-gRNAs) loci to analyze the F2 progeny of F1 females with identical genetic content but different inheritance of the constructs from the F0 parents: (**A**) Cas9 from the F0 male and gRNAs from the F0 female, (**B**) Cas9 from the F0 female and gRNAs from the F0 male, (**C**) both Cas9 and gRNAs from the F0 male, and (**D**) both Cas9 and gRNAs from the F0 female. (**E**) Analysis of the F2 inheritance rates of the fluorescent markers for all cross scheme combinations (**A-D**) using Cas9 constructs driven by the *vasa* promoter. Strong super-Mendelian inheritance is seen for all conditions except when both Cas9 and the gRNAs are inherited from the F0 female (**D**). Allelic conversion events are indicated by the red and green triangles in the F1 females of every cross scheme. Values for the inheritance average (black bar), standard deviation, number of samples (n) and total number of flies scored in each experiment are represented over the graph in line with the respective data.

### X-chromosome tGD uncovers maternal effect on inheritance

Previous studies reported that the inheritance of a gene drive from the female germline could lead to the generation of early embryogenesis mutations^5^, which occur when cleaved-allele repair results in small insertion/deletions (indels) at the cut site instead of allelic conversion, through alternative repair pathway such as the non-homologous end-joining (NHEJ) one^26^. Indel alleles therefore represent an obstacle for CRISPR gene-drive propagation in subsequent generations, as such alleles would be resistant to the gene-drive action, preventing its spread^2,5,16,17^. These resistant alleles prevent efficient gene drive in female-germline inheritance, most likely due to Cas9 deposition in the egg^4,5^. We therefore performed the reciprocal cross using our tGD(y,w) approach by collecting F0 *vasa*-Cas9 females and gRNA-carrying males (**Fig. 2B**). In the F2 analysis, we observed similar inheritance rates to the previous tGD analysis of 92% (Cas9-Red) and 97% (gRNAs-Green) (**Fig. 2E, Supplementary Data 2**), indicating no maternal inheritance effect is observed when Cas9 in inherited by the mother in this tGD arrangement like was observed in previous full-GD approaches^8,9,16^.

Since the two CRISPR components of our tGD are inherited separately, we could use this system to test whether both components had to be simultaneously deposited in the egg to observe the maternal effect. We generated a homozygous line carrying both elements on the same chromosome to analyze the allelic conversion efficiency for co-inheritance. As before, we analyzed the F2 progeny of cross schemes in which the coupled Cas9/gRNA elements were inherited together from either the F0 male (**Fig. 2C**) or female (**Fig. 2D**) in a condition that would mimic a full-GD scenario. Here, the F2 progeny from F0 male inheritance had inheritance rates of 91% (Cas9) and 95% (gRNA) (**Fig. 2E**). In contrast, the inheritance rates from the F0 female were 53% (Cas9) and 67% (gRNAs) (**Fig. 2E**), suggesting that a strong maternal effect on a gene drive is generated only when the two elements are inherited together from a female germline. These results encourage new gene drive designs that delay Cas9 and/or gRNA functioning in the embryo to avoid resistant allele formation^21^.

Next, to test if our tGD(y,w) would perform equally in terms of inheritance rates and maternal effects with a different promoter, we cloned the *nanos* gene regulatory region into our Cas9 construct and inserted it at the same location (*yellow*) to generate the *nanos*-Cas9 line (**Supplementary Fig. 1**). Performing the same cross schemes to combine the new *nanos*-Cas9-Red with the *y*-[*y1,w2*] line (**Fig. 2A-D**), a similar pattern was seen as for the *vasa* promoter, with comparable inheritance rates for separated transgenes in the F0 crosses and strong resistant allele formation when the combined Cas9/gRNA complex came from F0 females (**Supplementary Fig. 3, Supplementary Data 2**). An additional noteworthy result from this experiment was that our tGD(*y,w*) driven by *nanos* alleviated the inheritance differences between the Cas9 and gRNAs of our previous vasa-driven tGD(*y,w*) (**Fig. 2E; Supplementary Fig. 3**). These results reinforce that inheriting a preloaded Cas9-gRNA complex through the mother is an obstacle to gene drive spreading, which did not occur when the elements are inherited separately in our system.

### Gene drive process reveals predictable resistant allele formation

To better understand the maternal effect on gene-drive inheritance, we used males recovered from the F2 generation of the simultaneous-inheritance tGD(*y,w*) crosses that carried resistant alleles; since males have only one X chromosome, which is inherited from the F1 female, they are suitable for phenotypic isolation and molecular characterization of non-conversion events (resistant alleles) that occurred in each single F1 female. We analyzed resistant alleles in F2 males, from various experimental conditions, selected from 57 (*white*) and 60 (*yellow*) independent F1 female germlines, for a total of 242 and 225 flies sequenced per locus, respectively (**Fig. 3A,B, Supplementary Fig. 4**). Interestingly, for both *w* and *y*, we recovered three resistant alleles that occurred repeatedly in the germline of independent F1 females and that covered the majority of all sequenced flies (named wA, wB, wC and yA, yB, yC; **Fig. 3A,B, Supplementary Fig. 4**). Additionally, we observed fewer unique occurrences in *white* than in *yellow*, 18/242 (7%) and 43/225 (19%), respectively (**Fig. 3A,B**; light gray), highlighting the importance of characterizing the range of possible indels at genomic locations chosen for field gene-drive applications. Additionally, we analyzed the frequency of resistant mutations generated from conditions meant to resemble full-GD situations, with both elements inherited simultaneously (**Fig. 2C,D**), and observed that the ratios of recovered resistant alleles and their quality can vary drastically (**Supplementary Fig. 5**).

**Figure 3.**
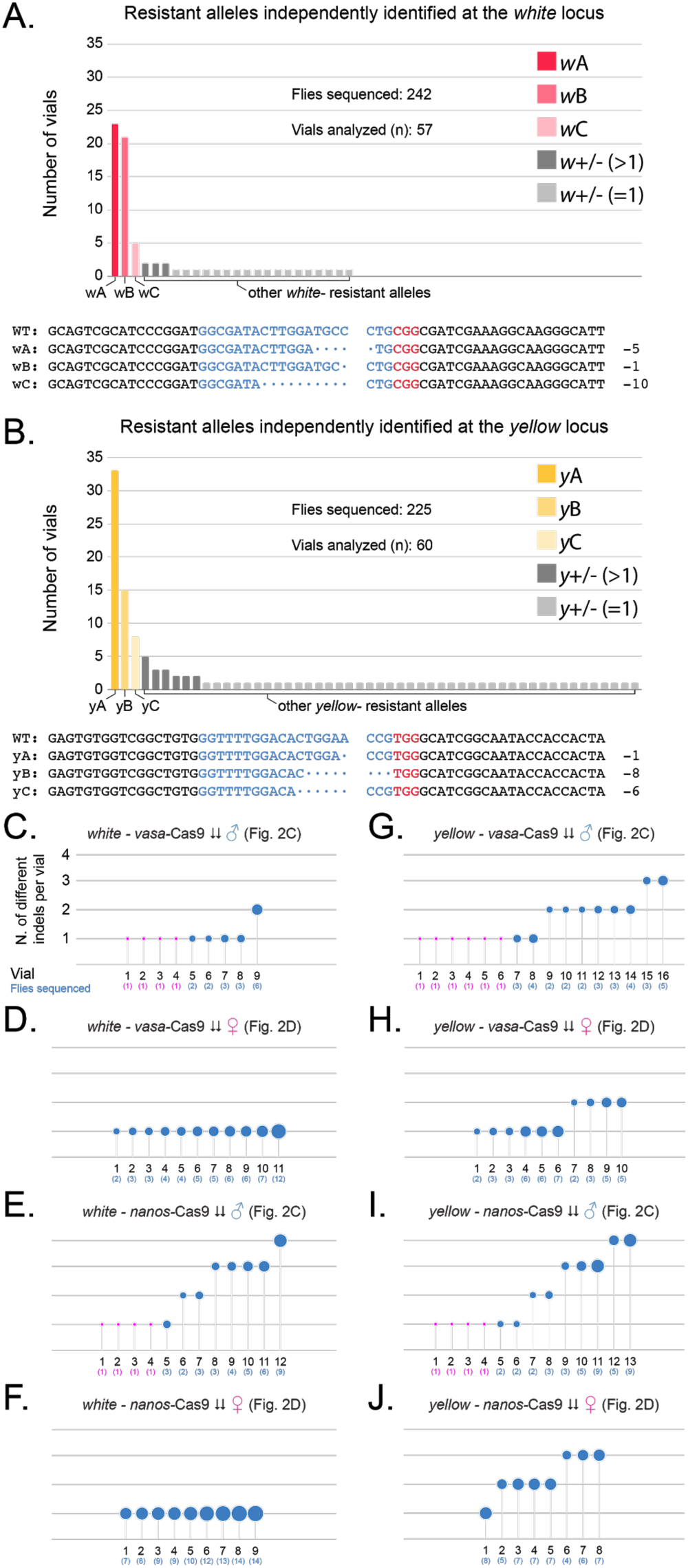
Analysis of the resistant alleles from the tGD(*y,w*) arrangement. (**A, B**) Graphs represent the independent generation of specific indel mutations generated in the experiments carried out in Fig. 2 and Fig. S3 when allelic conversion failed at the (**A**) *white* and (**B**) *yellow* loci. At both loci, we observe three repeatedly isolated indels that are colored with different shades of red for the *white* locus (wA, wB, wC) and yellow for the *yellow* locus (yA, yB, yC), the sequence of which is reported under each graph indicating with dots the missing bases compared to the wild-type sequence, split at the expected cut site. Additional indels recovered more than once are colored in dark grey, and those recovered only once are colored in light gray. (**C-J**) Each panel depicts the number of different indels recovered (y-axis) in ascending order for each F1 female (numbered on x-axis) for which the F2 male progeny was sampled. Bubble size is indicative of the number of flies analyzed for each specific F1 female, reported under the vial number in parenthesis. For F1 females producing only one F2 male with an indel, and therefore only one male was sampled, the dot is represented in magenta.

Lastly, in an attempt to clarify at what time point during development the allelic conversion process occurs, we also analyzed the number of different resistant alleles recovered in the F2 progeny of single F1 females, and in all cases, we detected a range of 1–4 different resistant alleles per vial (**Fig. 3C-J**). When the Cas9 and gRNAs constructs were inherited from the F0 female we recovered, at the *white* locus, only one resistant allele per vial analyzed (**Fig. 3D, F**), suggesting that these indels are generated as early as fertilization (or zygote), consistent with the average inheritance of ∼50% observed (*vasa* **Fig. 2D, E** and *nanos* **Supplementary Fig. 3**). This trend was not observed for the *yellow* locus, which showed up to three different alleles in the same conditions, suggesting that the promoter used to express the gRNAs or the gRNA itself results in lower efficiency of cutting at the *yellow* locus than the equivalent for *white* (**Fig. 3H, J**).

Regarding the male inheritance, we observe 1-4 indels generated from each F1 female germline for either promoter and locus analyzed **(Fig. 3C, E, G, I)**. This fact, combined with the high inheritance observed in these experiments suggests a stochastic generation of the resistant alleles during late germline development, after pole-cells formation (**Fig. 4A**). Additionally, in this condition, adult F1 females display w-mosaic eyes due to leakiness of the *vasa* and *nanos* promoters in somatic tissue, suggesting that the somatic tissue was not edited during early embryogenesis (**Fig. 4A**). Conversely, in female inheritance conditions, the indels seem to be generated in the syncytial blastoderm embryo before cellularization of the ∼3 pole cells that have been estimated to be set aside for the germline at the 128 cell stage^27^. In fact, at the *white* locus, we observe only 1 indel per female germline, which combined with the ∼50% inheritance measured for the gRNA transgene and the fully *w-* F1 female phenotype, suggest that Cas9 action occurs efficiently at the zygote stage (**Fig. 4B**). For *yellow* we instead observe 1-3 indels generated per F1 female, suggesting that Cas9 cutting happens later or less efficiently before pole-cells formation. In the case of *yellow*, our data is also consistent with few nuclei escaping Cas9 action in the blastoderm embryo and being converted later during germline development as we observe super-Mendelian inheritance in some of the F1 females (**Fig. 4C**; **Fig. 2D, E**; **Supplementary Fig. 3**).

**Figure 4.**
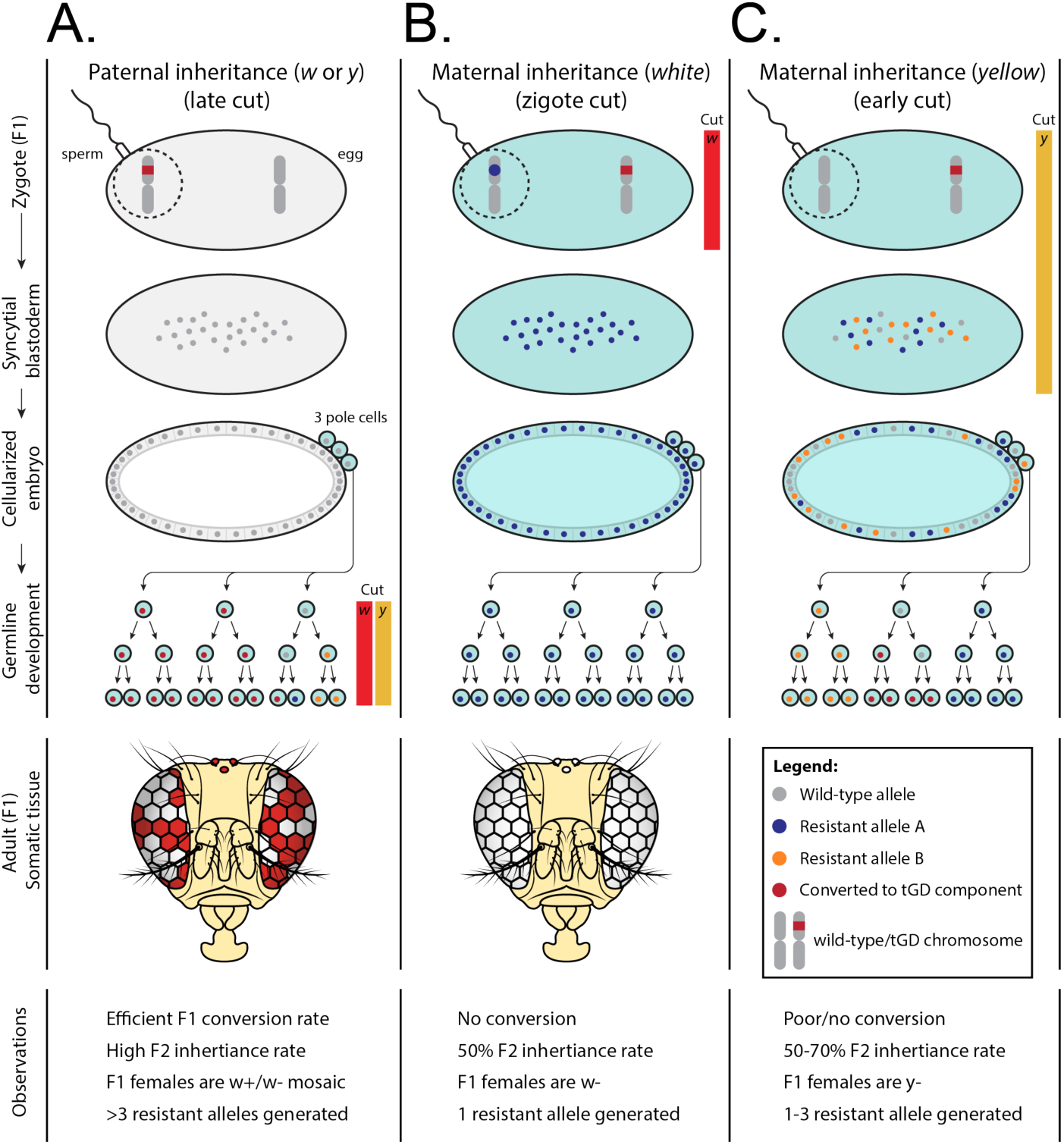
Model of tGD transgenes behavior in males and females. **(A-C)** Schematic representation of the different scenarios observed in our tGD experiments when both elements (Cas9 and gRNA transgenes) are inherited together. **(A)** When the tGD chromosome (labelled in red) comes from the dad (sperm) any Cas9 activity was detected from zygote to cellularized embryo. The high F2 inheritance rates in these conditions suggest that Cas9 action (red and yellow bars) is restricted to the germline development phase for the *white* and *yellow* genes, allowing efficient conversion (red dots) in this stage. Since the conversion process would occur after 3 pole cell formation, this would be consistent with the 1-4 resistant alleles range (blue and orange dots) captured from F1 germline females analyzed. **(B-C)** We detected different outputs for both *yellow* and *white* loci when the tGD chromosome was inherited from the mother; **(B)** Cas9 cut (red bar) seems to occur at the zygote stage for the *white* locus since we only observe one resistant allele per each F1 germline female that was analyzed. Consistently, this early cut event leads to 50% inheritance of our transgene without any possible conversion event. **(C)** The *yellow* locus edition by Cas9 (yellow bar) seems to be delayed based on the super-Mendelian inheritance observed in the scored F2 flies from some F1 germline females. In this case, Cas9 activity and the conversion process could occur in the zygote or syncytial blastoderm stage, and before pole cells establishment. This is reinforced by the presence of 1-3 resistant alleles range (blue and orange dots).

While our data is not conclusive, it strongly supports the hypothesis that in all the cases of female inheritance analyzed and previous reports using comparable reagents^1,5,16,17^, the Cas9-mediated cleavage leading to either allelic conversion or the resistant alleles is performed during early embryogenesis, before germline formation. Importantly, our system allowed us to evaluate, in the same animal, the simultaneous action of two gRNAs and identify how they differ in the maternal effect on super-Mendelian inheritance (**Fig. 4**).

### A chemo-genetic technology applied to tGD for controlling activation in the adult germline

Since our above-presented studies on female germline resistance suggest the Cas9-driven cuts could happen as early as the zygote stage, posing a potential problem to gene drive applications, we wondered how allelic conversion would perform when solely restricted to the adult germline. No published gene-drive work has thus far been able to precisely evaluate or modulate the timing of allelic conversion such that we could directly measure this.

We recently developed a small–molecule-controlled system for use in active genetics approaches including CRISPR-based gene-drive systems^28^. Briefly, we fused *Escherichia coli* dihydrofolate reductase (DHFR) domains to *Sp*Cas9 to promote its rapid proteasomal degradation in the absence of the stabilizing small molecule trimethoprim (TMP)^29,30^. We showed that TMP addition to the fruit fly diet stabilized our modified *Sp*Cas9 (DD2-Cas9) and allowed super-Mendelian inheritance control of a CopyCat active genetic element^28^.

Here, we first used a comparable DD2-Cas9 line and showed that mentioned drug-regulated system could be applied to the tGD(y,w) for controlling its super-Mendelian inheritance (**Supplementary Fig. 6; Supplementary Data 3**). Next, we used the TMP regulation in our tGD system and were able to activate Cas9 only in the adult female germline, showing that super-Mendelian inheritance can be achieved when the gene drive process is restricted to this tissue (**Supplementary Fig. 6; Supplementary Data 3**), although resistant alleles were also detected (**Supplementary Fig. 7**).

This approach opens a new avenue for restricting Cas9 activity to a proper window when HDR is favored, perhaps representing a way to bypass the maternal effect. Additionally, future developments of this technology could bias inheritance in, for example, a spatial fashion, such as by city, through the addition of the small molecule to urban water reservoirs, therefore controlling the spread of a gene drive into a circumscribed locale.

### Impaired homology asymmetrically affects drive efficiency

Recent research has raised concerns that natural polymorphisms could also hamper gene-drive spread in a population^31^, and a proposed strategy to increase drive efficiency and work around resistant alleles or polymorphisms aims to use multiple gRNAs to ensure cutting and lower the chances of an indel^17,32^. However, as it is intrinsically impossible to have perfect homology to all DNA ends generated when using multiple gRNAs, we wondered to what extent this homology discordance between the cleaved chromosome and the allele to be propagated would affect the efficiency of a gene drive. We reasoned that our tGD system would be an ideal tool to test this hypothesis, as we would be able to impair homology on the gRNA construct while keeping the untouched *vasa-*Cas9 element as an internal control.

For this purpose, we generated three modified versions of our *w-*[*y1,w2*] line that varied the location of a 20 bp lack in sequence homology: i) the first lacked 20 bp of homology on each side to generate the *w-*[*y1,w2*]-Short-HAs line (both sides impaired); ii) the second lacked 20 bp on the protospacer adjacent motif (PAM), which is an essential DNA-homing sequence for CRISPR function, side of the gRNA (PAM-proximal); and iii) the third lacked 20 bp on the side distal from the PAM (PAM-distal) (**Fig. 5, Supplementary Fig. 1**). We performed the same cross scheme as in **Fig. 2A** by combining the *vasa*-Cas9 line with each of the three lines bearing impaired homology and scoring the F2 progeny for inheritance rates.

**Figure 5.**
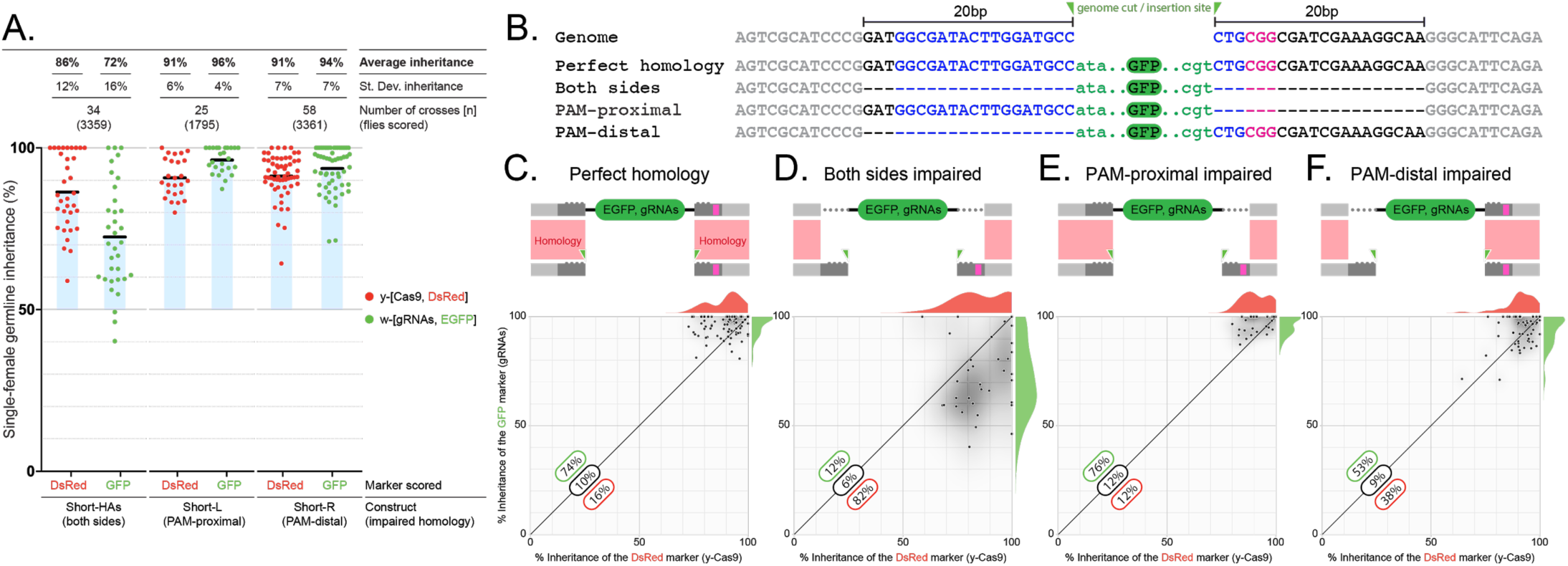
Impaired homology arms affect drive efficiency. (**A**) Inheritance rates of the three modified *w-*[*y1,w2*] gRNA lines (combined with *vasa*-Cas9 inserted in *yellow* gene) in which the homology (**B**) was impaired by removing 20 bp of sequence on both sides of the transgene (both sides), only on the PAM-proximal side (PAM proximal), or on the PAM-distal side (PAM distal). (**B**) Sequence representation of genome and transgene used, 20 base pairs impaired homology sequences indicated in black, blue and magenta; blue indicates the gRNA target, magenta the PAM sequence and green triangles highlight the gRNA driven cut location and gRNA-GFP transgene insertion site. The transgene sequence containing GFP is indicated in green. Dashes highlight the missing sequences in the impaired homology constructs. **(C-F)** A schematic of each construct showing how the homology is impaired during the repair process. The bottom portion of the panel contains a 2D-correlation plot in which the F2 inheritance of the DsRed-marked construct (Cas9 inserted in *yellow*) is represented on the x-axis, while the inheritance of the GFP-marked gRNA cassette is on the y-axis. The individual data points are overlaid on a 2D-density plot of the distribution, and a 1D-distribution of the data on each axis is also shown on the top and right of each graph. (**C**) Control: the 2D-correlation graph represents the data previously shown in **Fig. 2B** with perfect homology in both transgenes, representing the baseline. It is possible to observe how the gRNA construct performs overall better than the DsRed one (i.e., most dots fall over the diagonal line). (**D**) Homology impaired on both sides of the gRNA transgene dramatically reduces allelic conversion of the GFP-marked construct. (**E**) Impaired homology on the PAM-proximal side behaves similarly to the control. (**F**) Impaired homology on the PAM-distal side seems to be slightly affected in terms of GFP inheritance, as individual data points are distributed almost equally on both sides of the diagonal. (**C-E**) Quantification of the percentage of data points falling over (green), on (black) and under (red) the midline is overlays on each graph.

When both homology arms were impaired, we observed a significantly lower gRNA transgene inheritance, average of 72% (**Fig. 5A**, first condition; **Supplementary Data 4**), than the previously observed 96% with perfect homology (**Fig. 2E**, first condition). When we instead tested the PAM-proximal and PAM-distal lines, we did not observe any significant decrease, suggesting that imperfect homology can be tolerated as long as both sides are not impaired simultaneously (**Fig. 5A**; **Supplementary Data 4**). Importantly, our internal control Cas9 transgene averaged ∼90% inheritance rates in all conditions. Interestingly, the line with both sides impaired displayed a slightly lower inheritance for the Cas9 transgene and a significantly higher standard deviation when compared with the other conditions (**Fig. 5A; Supplementary Data 4**). These results also indicate that gene-drive applications with inherent imperfect homology are feasible, though the nature of the homology should be considered when designing multiplexed gRNA strategies.

When comparing plots showing the correlation between the inheritance rates for allelic conversion at the two loci, Cas9-red on the Y-axis and gRNA-Green on the X-axis (**Fig. 5C-F**), we observe that when the homology is lacking on the PAM-proximal side, the dots present a similar pattern as for the perfect homology construct, with the majority of dots located just over the diagonal (**Fig. 5C**). In contrast, it seems that there is an equivalent distribution of dots on both sides of the diagonal when the homology is lacking on the PAM-distal side (**Fig. 5F**). It has been shown that Cas9 can remain bound to the DNA after cleavage for an extended time^33–35^ and it preferentially releases the PAM-distal, non-target strand^36^. Our data suggests that this property might result in an asymmetrical influence of the bound Cas9/gRNA complex on the HDR process, and that perhaps the released PAM-distal, non-target strand may promote efficient HDR. This property might be harnessed for increasing HDR in systems that currently display poor efficiency, such as human somatic cells in therapeutic efforts^37^.

### Mathematical modeling predicts tGD alleles would collectively spread to a higher frequency than full gene-drive alleles

To predict the extent of tGD spread compared to full-GDs and determine tGD suitability for population replacement in the field, we performed mathematical modeling. Four tGD systems were considered: i) components linked on an autosome (tGD), ii) components unlinked at two autosomal loci (tGDc), iii) components linked on the X chromosome (tGDX), and iv) components unlinked on the X chromosome (tGDXc). We also considered full-GD systems: i) at an autosomal locus (full-GD), and ii) at an X-chromosome locus (full-GDX). For the purpose of model exploration, we estimated ballpark parameters for each system: i) cleavage frequency of 100%, ii) allelic conversion efficiency of 50–100%, and iii) no fitness costs associated with the Cas9 or gRNA alleles. All resistant alleles were assumed to be in-frame/cost-free. We modeled releases of *Aedes aegypti*, the mosquito vector of dengue, Zika, and chikungunya viruses, and simulated five weekly releases of 100 adult males homozygous for each system into a population with an equilibrium size of 10,000 adults. Model predictions were computed using 50 realizations of the stochastic implementation of the MGDrivE simulation framework^38^.

Exploratory results for these ballpark parameter estimates suggested that the tGDc system, spread across two loci, performs very similarly to a full-GD with some potentially beneficial qualities. At high allelic conversion efficiencies (90–100%), both systems spread at similar speeds; as the allelic conversion efficiency declined (50–90%), the full-GD spread slightly quicker than the tGDc system (**Fig. 6A-B**). Resistant alleles accumulated to similar overall proportions for both systems (**Fig. 6A**), though because the tGDc system is spread across two loci, a higher proportion of individuals had at least one copy of a transgene at equilibrium (for allelic conversion efficiencies <100%) (**Fig. 6C-D**), with almost all individuals having at least one copy of a transgene at equilibrium for allelic conversion efficiencies of 90–100%. This could be advantageous for population replacement strategies where a disease-refractory cassette could be linked to both the Cas9 and gRNA components of the tGDc system.

**Figure 6.**
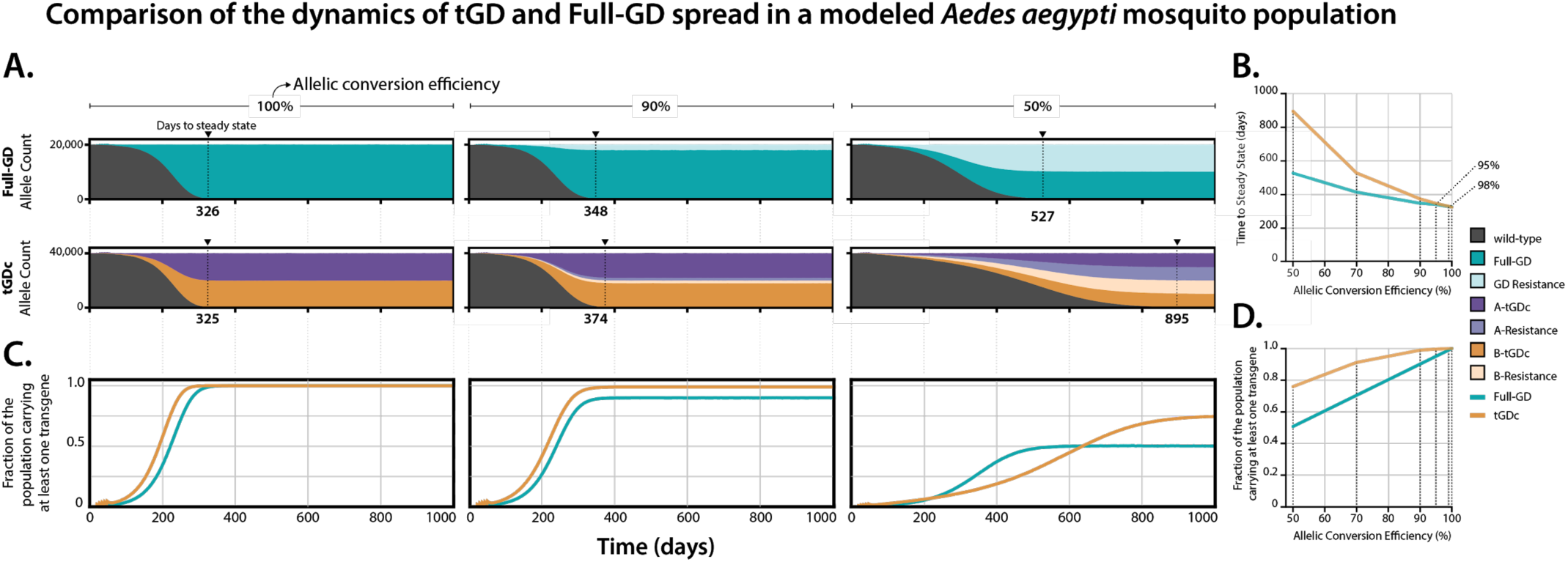
Mathematical modelling of full-drive and tGD spread through *Ae. aegypti* populations. Model predictions for releases of *Ae. aegypti* mosquitoes homozygous for the tGD and full-drive systems, parameterized with ballpark estimates: i) a cleavage frequency of 100% in females and males, ii) an allelic conversion efficiency given a cleavage of 50–100% in females and males, and iii) no fitness costs associated with the Cas9 or gRNA alleles. All resistant alleles are assumed to be in-frame/cost-free. Five weekly releases are simulated, consisting of 100 adult males homozygous for each system into a population having an equilibrium size of 10,000 adults. Model predictions were computed using 50 realizations of the stochastic implementation of the MGDrivE simulation framework^36^. **(A)** Stacked allele counts over time for the full drive and tGD systems for allelic conversion efficiencies of 100%, 90%, and 50%. **(B)** Allelic conversion efficiency to steady state plotted against time for the full drive (turquoise) and tGD (yellow) systems. **(C)** Fraction of the population carrying at least one transgene over time for the full drive (turquoise) and tGD (yellow) systems for allelic conversion efficiencies of 100%, 90%, and 50%. **(D)** Allelic conversion efficiency plotted against fraction of the population carrying at least one transgene for the full drive (turquoise) and tGD (yellow) systems.

Within the tGD systems, having the components on autosomal loci seems to be the most effective design based on this exploratory modeling exercise. Autosomal systems spread faster than X-linked systems due to their ability to drive in both sexes (**Supplementary Fig. 8A-B**). Interestingly, the linked tGD system spread slightly faster than the unlinked tGDc system at moderate-to-low allelic conversion efficiencies (∼50%), though this difference was modest and unnoticeable for higher allelic conversion frequencies (90–100%) (**Supplementary Fig. 8C**). Autosomal systems also result in a higher proportion of individuals with at least one copy of the transgene at equilibrium (**Supplementary Fig. 8C-D**). While these results are preliminary, neglecting fitness costs and detailed ecological considerations, they suggest potential benefits of the tGD system in the field that warrant further investigation.

## CONCLUSION

The tGD described herein splits the two genetic elements required for a gene drive into two separate locations that, when combined through genetic crossing, displayed super-Mendelian inheritance at three different loci (*e, y*, and *w*). We provide evidence for the advantages of a bipartite tGD system which allows flexible mix and match of gene-drive components to study key drive parameters and optimize overall drive efficiency before any field application. As opposed to a full-GD that propagates both Cas9 and gRNA as a single unit lacking intrinsic malleability to study their behavior, we show that the tGD is amenable to combinatorial optimization since the one element can be independently modified to study its effect on the overall gene drive efficiency.

Addressing one of the most pressing issues in the genetic-engineering community, our tGD approach also increases laboratory safety practices, since it greatly reduces potential spreading in case of accidental escape of the laboratory animals, as the Cas9 and gRNA elements are kept as different lines that are combined only during experimentation^15^.

Importantly, a mathematical simulation suggests significant advantages of using tGD over full-GD technologies for population modification. Due to its bi-component nature, the tGD can lead to a higher number of beneficial transgenes in a population at equilibrium (eg.: anti-malarial gene)^39^. Indeed, our work paves the way for safer gene-drive research and provides a quicker and more systematic gene-drive optimization strategy to help move these technologies to mosquitoes and other insect pests.

## ACKNOWLEDGEMENTS

We thank Bill McGinnis, Steve Wasserman, Mike Perry, Kaycie Butler and members of the Gantz laboratory for comments and edits on the manuscript. We thank Emily Bulger and Shannon Xu for experimental contribution on generating reagents. Research reported in this manuscript was supported by the University of California, San Diego, Department of Biological Sciences, by the Office of the Director of the National Institutes of Health under award number DP5OD023098 and by a DARPA Safe Genes Program Grant (Brdi N66001-17-2-4055). A Paul G. Allen Frontiers Group Distinguished Investigators Award supported E.B., a gift from the Tata Trusts of India to TIGS-UCSD supported X.F., a DARPA Safe Genes Program Grant (HR0011-17-2-0047) supported J.B. and J.M.M., and funds from the UC Irvine Malaria Initiative supported H.M.S.C. and J.M.M. The fruit fly drawing used in Figure 4 was adapted from original artwork by Madboy74 [CC BY-SA 4.0 (https://creativecommons.org/licenses/by-sa/4.0)].

## AUTHOR CONTRIBUTIONS

E.B. and V.M.G conceived the project. V.L.D.A. and V.M.G contributed to the design of the experiments V.L.D.A., V.M.G, A.L.B., and X.F. performed the experiments and contributed to the collection and analysis of data. H.M.S.C., J.B., and J.M.M. designed and performed the mathematical modeling experiments. V.L.D.A., J.M.M., and V.M.G. wrote the manuscript. All authors edited the manuscript.

## METHODS

### Fly rearing and maintenance for experiments

Fly stocks were raised at 18°C with a 12/12 hour day/night cycle on regular cornmeal medium. Experimental flies were grown at 25°C with a 12/12 hour day/night cycle. For TMP experiments, we used Formula 4-24 Instant Drosophila Food (Carolina Biological Supply Company, Cat. # 173214). After weighing 1 gram of food per tube, we reconstituted it by adding 3ml of water with DMSO (Fisher Scientific, Cat. #D128) or water containing different concentrations of TMP (Oakwood Chemical, Cat. # 036441) dissolved in DMSO (Fisher Scientific, Cat. # D128). Flies were anesthetized using CO2 to select individuals for crossing and phenotyping, and were scored by tracking fluorescent markers with a Leica M165 FC Stereo microscope with fluorescence. We used the DsRed and EGFP (referred to in the main text as GFP) markers as evidence for successful conversion. For the yellow body phenotype, we did not track mosaicism. For the white eye phenotype, we scored white, red, or mosaic eyes (see **Supplementary Data 1-4**). All crosses were performed following the Institutional Biosafety Committee-approved protocol from University of California San Diego. Every gene-drive experiment is performed in a high-security Arthropod Containment Level 2 (ACL2) barrier facility in plastic vials that are autoclaved before being discarded. Additionally, this facility requires two levels of security card access.

### Plasmid construction

Standard molecular biology techniques were used to generate all constructs analyzed in this work. Final sequence information of constructs used in this work will be available on NCBI before publication and accession numbers will be provided.

### Transgenic line generation and genotyping

Constructs inserted at the *ebony, yellow*, and *white* loci were co-injected with a Cas9 expressing plasmid (Addgene #46294) and a pCFD3 plasmid (Addgene #49410) expressing previously validated gRNA-*e1*^*1*^, gRNA-*y1*^*2,3*^, or gRNA-*w2*^*3*^, respectively. Constructs used for transgenesis are outlined in **Supplementary Fig. 1**. We marked the Cas9 and gRNA constructs with either a DsRed (Red) or an EGFP (Green) fluorescent reporter expressed in the eye using the 3xP3 promoter. All injections to generate transgenic flies were performed by BestGene, Inc. or Rainbow Transgenic Flies, Inc. All constructs were injected into an isogenized Oregon-R (OrR) strain from our laboratory to ensure a consistent background throughout all our experiments. After sending the constructs to the injection companies, we received 80–120 injected larvae. Once they hatched, we placed all G0 adults in different tubes (5–6 females crossed to 5–6 males). Then, G1 progeny were screened for positive flies with a specific fluorescent marker expressed in the eyes, which was indicative of successful transgene insertion. Flies positive for the marker were crossed individually to the same OrR flies we used for injection to make a homozygous stock in subsequent generations by identifying the *e, y* or *w* visible marker. Lastly, we sequenced each stock to confirm correct transgene integration.

### Molecular analysis of resistant alleles

For resistant allele sequence analysis we performed single fly DNA extractions using 50 ul of DNA extraction buffer (1M Tris, 0.5M EDTA, 5M NaCl, and Proteinase K [NEB, Cat. #P8107]). We squished individual flies for 1 minute, then placed them in a thermocycler for 30 minutes at 37°C, then 2 minutes at 95°C to inactivate the Proteinase K. Lastly, we added 200uL of water to dilute each sample to a final volume of 250uL. We used 1-5uL of each DNA extraction as a template for a 25uL PCR reaction. We performed PCRs covering the gRNA cute site for either the *yellow* or *white* locus in order to sequence the resistant allele present.

### Statistical analysis

We used GraphPad Prism 7 to perform all statistical analyses.

### Mathematical modeling

To model the expected performance of the trans-complementing gene drive system in populations of *Aedes aegypti*, the mosquito vector of dengue, chikungunya, and Zika viruses, we simulated release schemes for the trans-complementing system with: i) components linked on an autosome (tGD), ii) components unlinked at two autosomal loci (tGDc), iii) components linked on the X chromosome (tGDX), and iv) components unlinked at two loci on the X chromosome (tGDXc). We also compared the system to standard full gene drives at an autosomal locus (Full-GD), and at an X chromosome locus (Full-GDX). Releases were simulated consisting of 5 weekly releases of 100 adult males homozygous for each system using the MGDrivE simulation framework^4^ (https://marshalllab.github.io/MGDrivE/). This framework models the egg, larval, pupal, and adult mosquito life stages (both female and male adults are modeled) implementing a daily time step, overlapping generations, and a mating structure in which adult males mate throughout their lifetime while adult females mate once upon emergence, retaining the genetic material of the adult male with whom they mate for the duration of their adult lifespan. Density-independent mortality rates for the juvenile life stages are assumed to be identical and are chosen for consistency with the population growth rate in the absence of density-dependent mortality. Additional density-dependent mortality occurs at the larval stage, the form of which is taken from previous studies^5^. The inheritance patterns for the tGD, tGDc, tGDX, tGDXc, Full-GD and Full-GDX systems are modeled within the inheritance module of the MGDrivE framework^4^. We parameterized our trans-complementing and full gene drive models using ballpark parameter estimates for model exploration: i) a cleavage frequency of 100% in females and males, ii) a frequency of accurate homology-directed repair given cleavage of 50-100% in females and males, iii) no fitness costs associated with the Cas9 or gRNA alleles, and iv) all resistant alleles being in-frame/cost-free. We implemented the stochastic version of the MGDrivE framework to capture the randomness associated with low genotype frequencies and rare events such as resistant allele generation under some parameterizations. *Ae. aegypti* life history parameter values are listed below.

Table. Parameter values used in *Aedes aegypti* population model.

| Parameter | Symbol | Value | Reference |
| --- | --- | --- | --- |
| Egg production per female (day<sup>-1</sup>) | $\beta$ | 20 | <sup>6</sup> |
| Duration of egg stage (days) | $T_E$ | 5 | <sup>7</sup> |
| Duration of larval stage (days) | $T_L$ | 6 | <sup>7</sup> |
| Duration of pupa stage (days) | $T_P$ | 4 | <sup>7</sup> |
| Daily population growth rate (day<sup>-1</sup>) | $r$ | 1.175 | <sup>8</sup> |
| Daily mortality rate of adult stage (day<sup>-1</sup>) | $\mu_M$ | 0.090 | <sup>9,10</sup> |
| Adult female population size | $N$ | 10,000 | <sup>11</sup> |

## REFERENCES

1. Gantz, V. M. & Bier, E. Genome editing. The mutagenic chain reaction: a method for converting heterozygous to homozygous mutations. Science 348, 442–444 (2015).

2. Champer, J. et al. Novel CRISPR/Cas9 gene drive constructs reveal insights into mechanisms of resistance allele formation and drive efficiency in genetically diverse populations. PLoS Genet. 13, e1006796 (2017).

3. Kyrou, K. et al. A CRISPR–Cas9 gene drive targeting doublesex causes complete population suppression in caged Anopheles gambiae mosquitoes. Nat. Biotechnol. (2018). doi:10.1038/nbt.4245

4. Hammond, A. et al. A CRISPR-Cas9 gene drive system targeting female reproduction in the malaria mosquito vector Anopheles gambiae. Nat. Biotechnol. 34, 78–83 (2016).

5. Gantz, V. M. et al. Highly efficient Cas9-mediated gene drive for population modification of the malaria vector mosquito Anopheles stephensi. Proc. Natl. Acad. Sci. U. S. A. 112, E6736–43 (2015).

6. Noble, C., Adlam, B., Church, G. M., Esvelt, K. M. & Nowak, M. A. Current CRISPR gene drive systems are likely to be highly invasive in wild populations. Elife 7, (2018).

7. Marshall, J. & Akbari, O. Can CRISPR-based gene drive be confined in the wild? A question for molecular and population biology. (2017). doi:10.1101/173914

8. Courtier-Orgogozo, V., Morizot, B. & Boëte, C. Agricultural pest control with CRISPR-based gene drive: time for public debate. EMBO Rep. 18, 878–880 (2017).

9. McFarlane, G. R., Whitelaw, C. B. A. & Lillico, S. G. CRISPR-Based Gene Drives for Pest Control. Trends Biotechnol. 36, 130–133 (2018).

10. Grunwald, H. A. et al. Super-Mendelian inheritance mediated by CRISPR/Cas9 in the female mouse germline. (2018). doi:10.1101/362558

11. Esvelt, K. M., Smidler, A. L., Catteruccia, F. & Church, G. M. Concerning RNA-Guided Gene Drives for the Alteration of Wild Populations. (2014). doi:10.1101/007203

12. Adelman, Z. et al. Rules of the road for insect gene drive research and testing. Nat. Biotechnol. 35, 716–718 (2017).

13. James, A. A. Gene drive systems in mosquitoes: rules of the road. Trends Parasitol. 21, 64–67 (2005).

14. Committee on Gene Drive Research in Non-Human Organisms: Recommendations for Responsible Conduct, Board on Life Sciences, Division on Earth and Life Studies & National Academies of Sciences, Engineering, and Medicine. Gene Drives on the Horizon: Advancing Science, Navigating Uncertainty, and Aligning Research with Public Values. (National Academies Press (US), 2016).

15. Akbari, O. S. et al. BIOSAFETY. Safeguarding gene drive experiments in the laboratory. Science 349, 927–929 (2015).

16. Hammond, A. M. et al. The creation and selection of mutations resistant to a gene drive over multiple generations in the malaria mosquito. PLoS Genet. 13, e1007039 (2017).

17. Champer, J. et al. Reducing resistance allele formation in CRISPR gene drive. Proc. Natl. Acad. Sci. U. S. A. (2018). doi:10.1073/pnas.1720354115

18. DiCarlo, J. E., Chavez, A., Dietz, S. L., Esvelt, K. M. & Church, G. M. Safeguarding CRISPR-Cas9 gene drives in yeast. Nat. Biotechnol. 33, 1250–1255 (2015).

19. Gantz, V. M. & Bier, E. The dawn of active genetics. Bioessays 38, 50–63 (2016).

20. Xu, X.-R. S., Gantz, V. M., Siomava, N. & Bier, E. CRISPR/Cas9 and active genetics-based trans-species replacement of the endogenous -L2 CRM reveals unexpected complexity. Elife 6, (2017).

21. Champer, J. et al. Molecular safeguarding of CRISPR gene drive experiments. Elife 8, (2019).

22. Li, M. et al. Development of a Confinable Gene-Drive System in the Human Disease Vector, Aedes aegypti. doi:10.1101/645440

23. Wittkopp, P. J., True, J. R. & Carroll, S. B. Reciprocal functions of the Drosophila yellow and ebony proteins in the development and evolution of pigment patterns. Development 129, 1849–1858 (2002).

24. Ren, X. et al. Enhanced specificity and efficiency of the CRISPR/Cas9 system with optimized sgRNA parameters in Drosophila. Cell Rep. 9, 1151–1162 (2014).

25. Doench, J. G. et al. Rational design of highly active sgRNAs for CRISPR-Cas9-mediated gene inactivation. Nat. Biotechnol. 32, 1262–1267 (2014).

26. Chang, H. H. Y., Pannunzio, N. R., Adachi, N. & Lieber, M. R. Non-homologous DNA end joining and alternative pathways to double-strand break repair. Nat. Rev. Mol. Cell Biol. 18, 495–506 (2017).

27. Lindsley, D. L., Hardy, R. W., Ripoll, P. & Lindsley, D. Gonadal Mosaicism Induced by Chemical Treatment of Sperm in Drosophila melanogaster. Genetics 202, 157–174 (2016).

28. Lopez Del Amo, V. et al. Small-molecule control of super-Mendelian inheritance in gene drives. doi:10.1101/665620

29. Maji, B. et al. Multidimensional chemical control of CRISPR–Cas9. Nat. Chem. Biol. 13, 9–11 (2016).

30. Iwamoto, M., Björklund, T., Lundberg, C., Kirik, D. & Wandless, T. J. A general chemical method to regulate protein stability in the mammalian central nervous system. Chem. Biol. 17, 981–988 (2010).

31. Drury, D. W., Dapper, A. L., Siniard, D. J., Zentner, G. E. & Wade, M. J. CRISPR/Cas9 gene drives in genetically variable and nonrandomly mating wild populations. Sci Adv 3, e1601910 (2017).

32. Marshall, J. M., Buchman, A., Sánchez C, H. M. & Akbari, O. S. Overcoming evolved resistance to population-suppressing homing-based gene drives. Sci. Rep. 7, 3776 (2017).

33. Sternberg, S. H., Redding, S., Jinek, M., Greene, E. C. & Doudna, J. A. DNA interrogation by the CRISPR RNA-guided endonuclease Cas9. Nature 507, 62–67 (2014).

34. Shibata, M. et al. Real-space and real-time dynamics of CRISPR-Cas9 visualized by high-speed atomic force microscopy. Nat. Commun. 8, 1430 (2017).

35. Yang, M. et al. The Conformational Dynamics of Cas9 Governing DNA Cleavage Are Revealed by Single-Molecule FRET. Cell Rep. 22, 372–382 (2018).

36. Richardson, C. D., Ray, G. J., DeWitt, M. A., Curie, G. L. & Corn, J. E. Enhancing homology-directed genome editing by catalytically active and inactive CRISPR-Cas9 using asymmetric donor DNA. Nat. Biotechnol. 34, 339–344 (2016).

37. Orthwein, A. et al. A mechanism for the suppression of homologous recombination in G1 cells. Nature 528, 422–426 (2015).

38. Sánchez C., H. M., Wu, S. L., Bennett, J. B. & Marshall, J. M. MGDrivE: A modular simulation framework for the spread of gene drives through spatially-explicit mosquito populations. doi:10.1101/350488

39. Isaacs, A. T. et al. Transgenic Anopheles stephensi coexpressing single-chain antibodies resist Plasmodium falciparum development. Proc. Natl. Acad. Sci. U. S. A. 109, E1922–30 (2012).

## REFERENCES FOR THE METHODS SECTION

1. Port, F., Chen, H.-M., Lee, T. & Bullock, S. L. Optimized CRISPR/Cas tools for efficient germline and somatic genome engineering in Drosophila. Proc. Natl. Acad. Sci. U. S. A. 111, E2967–76 (2014).

2. Gantz, V. M. & Bier, E. Genome editing. The mutagenic chain reaction: a method for converting heterozygous to homozygous mutations. Science 348, 442–444 (2015).

3. Bassett, A. R., Tibbit, C., Ponting, C. P. & Liu, J.-L. Highly Efficient Targeted Mutagenesis of Drosophila with the CRISPR/Cas9 System. Cell Rep. 6, 1178–1179 (2014).

4. C. H. M. S., Sánchez C., H. M., Wu, S. L., Bennett J. B. & Marshall, J. M. MGDrivE: A modular simulation framework for the spread of gene drives through spatially-explicit mosquito populations. doi:10.1101/350488

5. Deredec, A., Godfray, H. C. J. & Burt, A. Requirements for effective malaria control with homing endonuclease genes. Proc. Natl. Acad. Sci. U. S. A. 108, E874–80 (2011).

6. Otero, M., Solari, H. G. & Schweigmann, N. A stochastic population dynamics model for Aedes aegypti: formulation and application to a city with temperate climate. Bull. Math. Biol. 68, 1945–1974 (2006).

7. Christophers, S. R. Aëdes Aegypti (L.), the Yellow Fever Mosquito: Its Life History, Bionomics, and Structure. (Cambridge University Press, 1960).

8. Simoy, M. I., Simoy, M. V. & Canziani, G. A. The effect of temperature on the population dynamics of Aedes aegypti. Ecol. Modell. 314, 100–110 (2015).

9. Focks, D. A., Haile, D. G., Daniels, E. & Mount, G. A. Dynamic life table model for Aedes aegypti (Diptera: Culicidae): analysis of the literature and model development. J. Med. Entomol. 30, 1003–1017 (1993).

10. Horsfall, W. R. Mosquitoes: their bionomics and relation to disease. (1972).

11. Carvalho, D. O. et al. Suppression of a Field Population of Aedes aegypti in Brazil by Sustained Release of Transgenic Male Mosquitoes. PLoS Negl. Trop. Dis. 9, e0003864 (2015).

