## Supplementary Information for "Split-gene drive system provides flexible application for safe laboratory investigation and potential field deployment"

#### Supplementary Figure 1

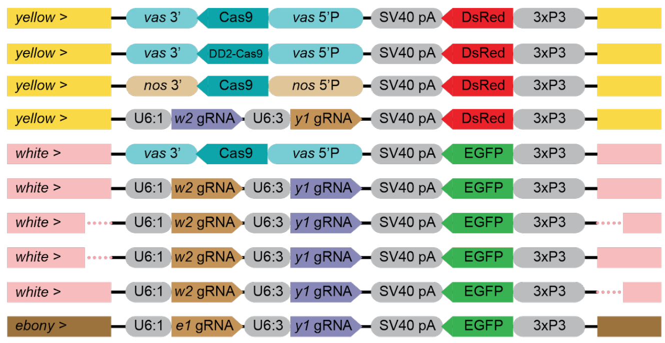


| Construct | Locus | Transgene | Used in figures | Generated |
|---|---|---|---|---|
| pVG182 | *yellow* | *vasa*-Cas9 | 1, 2, 5, Supp. 6 | *Lopez Del Amo et al. 2019* |
| pVG312 | *yellow* | *vasa*-DD2-Cas9 | Supp. 6 | *Lopez Del Amo et al. 2019* |
| pVG303 | *yellow* | *nanos*-Cas9 | Supp. 3 | This work |
| pVG186 | *yellow* | w2, y1 gRNAs | Supp. 2 | This work |
| pVG316 | *white* | *vasa*-Cas9 | Supp. 2 | This work |
| pVG185 | *white* | w2, y1 gRNAs | 2, 5, Supp. 3 | This work |
| pVG307 | *white* | Short-HAs (both sides) | 5 | This work |
| pVMG47 | *white* | Short-L (PAM-proximal) | 5 | This work |
| pVMG48 | *white* | Short-R (PAM-distal) | 5 | This work |
| pVG304 | *ebony* | e1, y1 gRNAs | 1 | This work |

**Supplementary Figure 1 - Constructs generated for transgenesis.** All *vasa* and *nanos* Cas9 lines contain the same *Sp*Cas9 sequence. Two Cas9 lines driven by *vasa* or *nanos* are inserted in *yellow* gene (pVG182 and pVG303 constructs, respectively), and flanked by specific yellow homology arms (yellow boxes) to allow allelic conversion of these transgenes when combined with gRNAs tandem. An additional Cas9 line driven by *vasa* promoter was instead inserted in *white* gene with specific homology arms to allow allelic conversion (construct pVG316). A modified version of pVG182 includes *Escherichia coli* dihydrofolate reductase (ecDHFR) domains that induce Cas9 degradation absence of trimethoprim (TMP). Two different tandem-gRNAs constructs were generated having either the *w2* and *y1* gRNAs or the *e1* and *y1* ones under the control of the U6:1 and U6:3 promoters respectively, inserted at different genomic locations (pVG186 in *yellow*, pVG185, pVG307, pVMG47 and pVMG48 in *white* and pVG304 in *ebony*). Fluorescent markers driven by the eye promoter 3xP3 were used to track each transgene. DsRed was used for transgenes inserted in *yellow* and EGFP was used for constructs targeting *white* or *ebony*. The table on the side summarizes this information including references to what figures in the manuscript use each construct. Constructs pVG128 and pVG312 were previously described [1].

**Supplementary Figure 2**

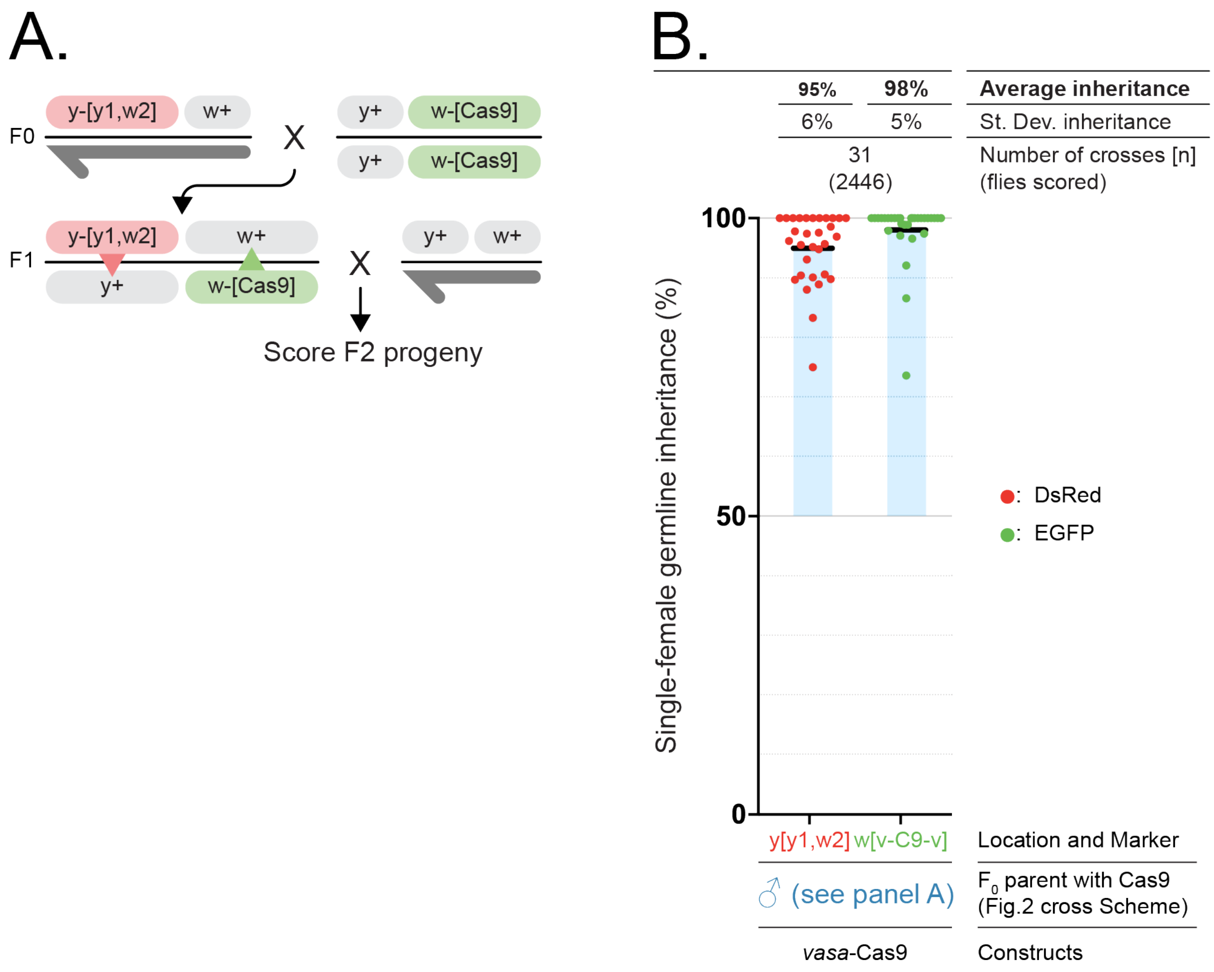


**Supplementary Figure 2 - The tGD(w,y) combination alleviates differences observed between Cas9 and gRNA transgenes inheritance observed in the tGD(y,w)**. The tGD(w,y) "swapped" version has the same Cas9 and gRNAs transgenes than in our tGD(*y,w*), but they are swapped in their insertion locus with the Cas9 inserted in *white* and the gRNAs in *yellow*. (**A**) Shows the genetic cross performed to analyze the F2 progeny. (B) Displays the data points in a plot highlighting the biased inheritance observed. Values for the inheritance average (black bar), standard deviation, number of samples (n) and total number of flies scored in each experiment are represented over the graph in line with the respective data.

**Supplementary Figure 3**

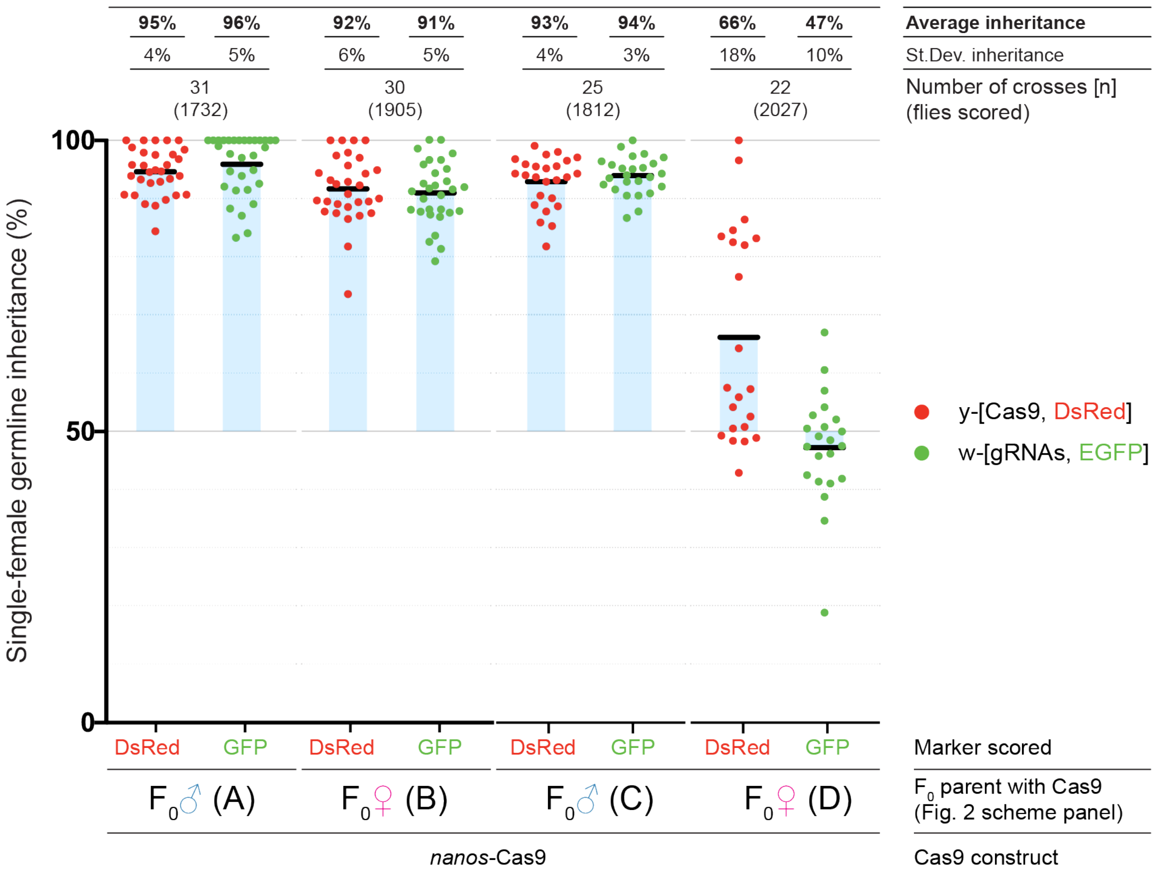


**Supplementary Figure 3 - tGD targeting the *yellow* and *white* loci driven by *nanos* promoter.** Analysis of the F2 inheritance rates of the fluorescent markers for the cross scheme combinations from **Fig. 2A-D** using a Cas9 construct driven by the *nanos* promoter. The inheritance observed in the F2 progeny of different F1 females are graphed: (**A**) Cas9 from the F0 male and gRNAs from the F0 female, (**B**) Cas9 from the F0 female and gRNAs from the F0 male, (**C**) both Cas9 and gRNAs from the F0 male, and (**D**) both Cas9 and gRNAs from the F0 female. The blue shading represents the deviation from the expected 50% “Mendelian” inheritance. Values for the inheritance average (black bar), standard deviation, number of samples (n) and total number of flies scored in each experiment are represented over the graph in line with the respective data.

### Supplementary Figure 4

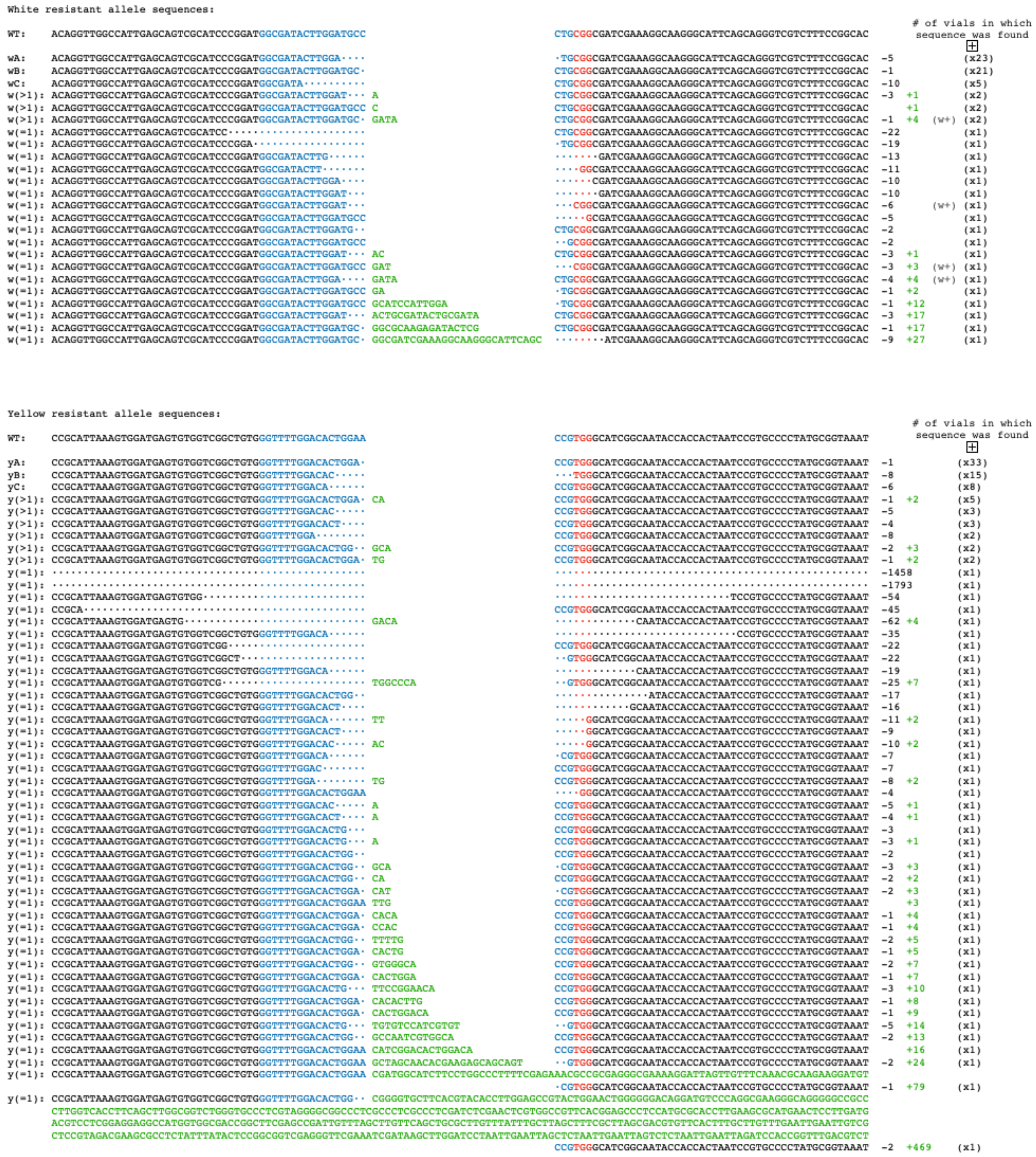


**Supplementary Figure 4 - Sequencing of resistant alleles generated in our tGD(y,w) experiments at both *white* and *yellow* loci.** These sequences were recovered by sequencing F2 males recovered from crosses performed in **Fig. 2** and **Supp. Fig. 3**. On top of each list, *white* (top) and *yellow* (bottom), the wild type (WT) sequence is represented. PAM sequence is shown in red, gRNA target sequence in blue, dots represent deleted nucleotides, green letters represent inserted nucleotides. On the left is reported the sequence nomenclature from **Fig. 3**. On the right, the number of deleted nucleotides in black, followed by number of inserted nucleotides in green, followed by the number of independent vials from which each sequence was recovered. For *white* if the analyzed fly displayed a wild-type phenotype it is followed by a *w+* label.

### Supplementary Figure 5

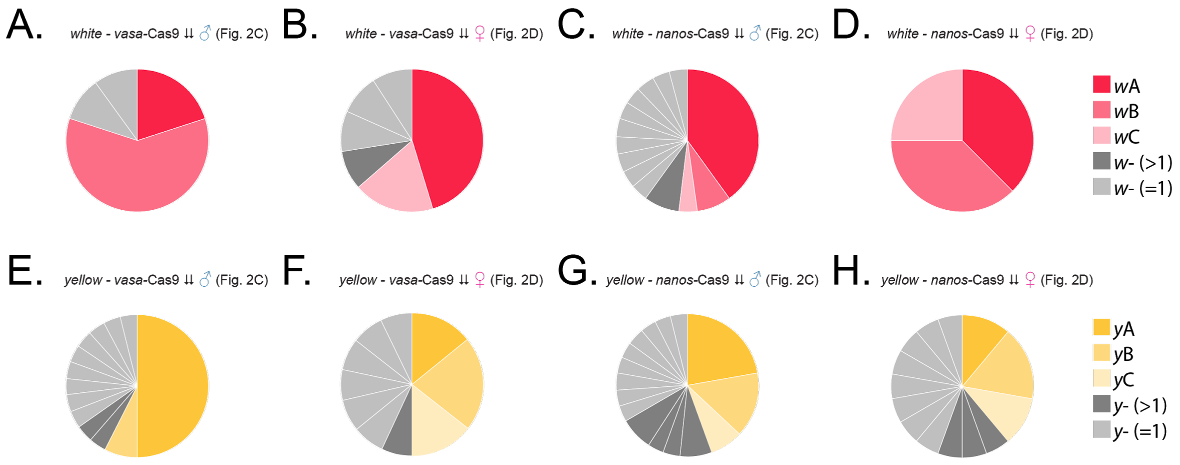


**Supplementary Figure 5 - Resistant allele distribution across tGD experiments using different cross schemes.** For the cross scheme in **Fig. 2C-D** we represent in pie charts the distribution of resistant alleles generated at the *white* (**A-D**) and *yellow* (**E-H**) loci when gene drive elements were inherited together and driven by either the *vasa* and *nanos* promoter. Each resistant allele displayed in the pie charts above has independently arisen in different vials. This graphs highlight that specific indel events are favored over others in different conditions. (**A**, **E**) *vasa*-Cas9 and gRNAs inherited from the F0 male. (**B**, **F**) *vasa*-Cas9 and gRNAs inherited from the F0 female. (**C**, **G**) *nanos*-Cas9 and gRNAs inherited from the F0 male. (**D**, **H**) *nanos*-Cas9 and gRNAs inherited from the F0 female. The color-code legend on the right corresponds to the overall abundance of indel observed in all our experiments, summarized in **Fig. 3**.

### Supplementary Figure 6

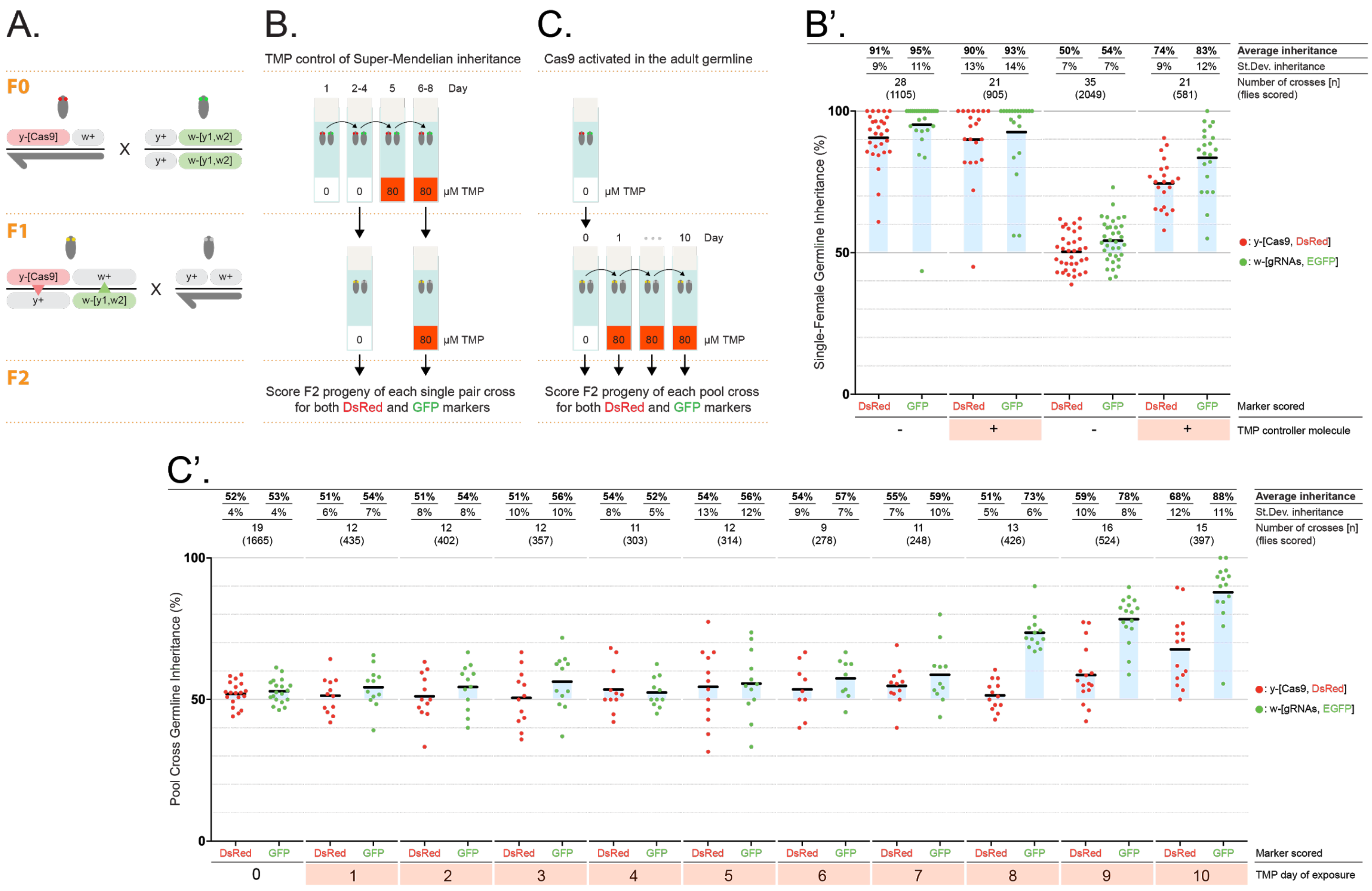


**Supplementary Figure 6 - Drug-inducible tGD system allows gene-drive control through a small-molecule feeding. (A)** Cross strategy performed to test drug-controlled activation of the tGD system. (**B**) Drug feeding strategy to test gene-drive activation during the entirety of fly development. F0 males carrying either the wild-type Cas9 or the drug-regulated version (DD2-Cas9), both inserted in *yellow* and driven by the *vasa* promoter, were crossed to F0 females carrying the tandem gRNA transgene in *white*. The F0 couple was kept on regular food for one day, allowed to lay eggs on a separate vial from day 2–4, kept for one day on a "conditioning" vial containing trimethoprim (TMP), and then passed onto a last vial for egg laying from day 6–8. Emerging F1 virgin females from the second (day 2–4) and fourth (day 6–8) vials were crossed to wild-type males on the same food condition to analyze the F1 germline transmission of the two transgenes by scoring the markers in the F2 progeny. This specific cross scheme allowed us to evaluate the transmission of the fluorescent markers through the germline of F1 sister females, minimizing potential differences due to genetic background. **(B')** Cas9 crosses are not affected by the presence of TMP, while DD2-Cas9 crosses display Mendelian inheritance of the markers in absence of TMP and super-Mendelian inheritance when TMP was added to the fly diet. **(C)** Drug feeding scheme to selectively activate allelic conversion in the adult germline. F0 cross was performed in regular food, and F1 virgin trans-heterozygous female progeny were collected and crossed first in small pools of 3–4 females to wild-type males on regular fly food. After that, each subsequent day the cross pool was passed onto a new fly tube containing 80 μM of TMP. (**C'**) The F2 progeny emerging from each tube were analyzed for the inheritance rates of the markers. An average inheritance of ~50% was observed in vials from days 0–7, and an increase in inheritance was seen for days 8–10, suggesting that gametes producing F2 offspring from days 0–7 had already undergone meiosis at the time of TMP exposure.

### Supplementary Figure 7

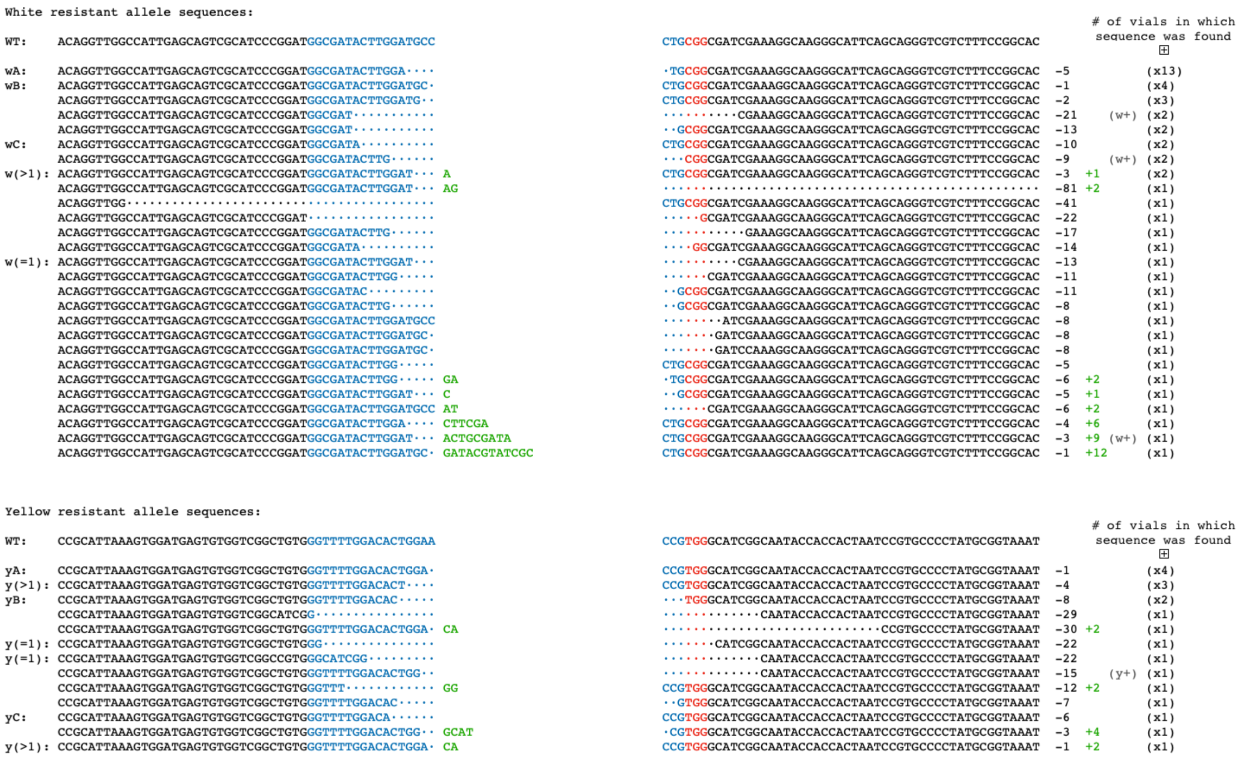


**Supplementary Figure 7 - Resistant allele sequences recovered from the drug-mediated activation of the tGD(y,w) in the adult germline.** These sequences were recovered by sequencing non-converted F2 males from **Supplementary Fig. 6** representing resistant alleles events generated in the adult germline of F1 females at the *white* (top) and *yellow* (bottom) loci. On top of each list the wild-type (WT) sequence is represented. PAM sequence is shown in red, gRNA target sequence in blue, dots represent deleted nucleotides, green letters represent inserted nucleotides. On the left is reported the sequence nomenclature from **Fig. 3**. On the right, the number of deleted nucleotides in black, followed by number of inserted nucleotides in green, followed by the number of independent vials from which each sequence was recovered. If the analyzed fly displayed a wild type phenotype it is marked with either *w+* or *y+*.

### Supplementary Figure 8

**Comparison of the dynamics of additional tGD and Full-GD versions as they spread in a modeled *Aedes aegypti* mosquito population**

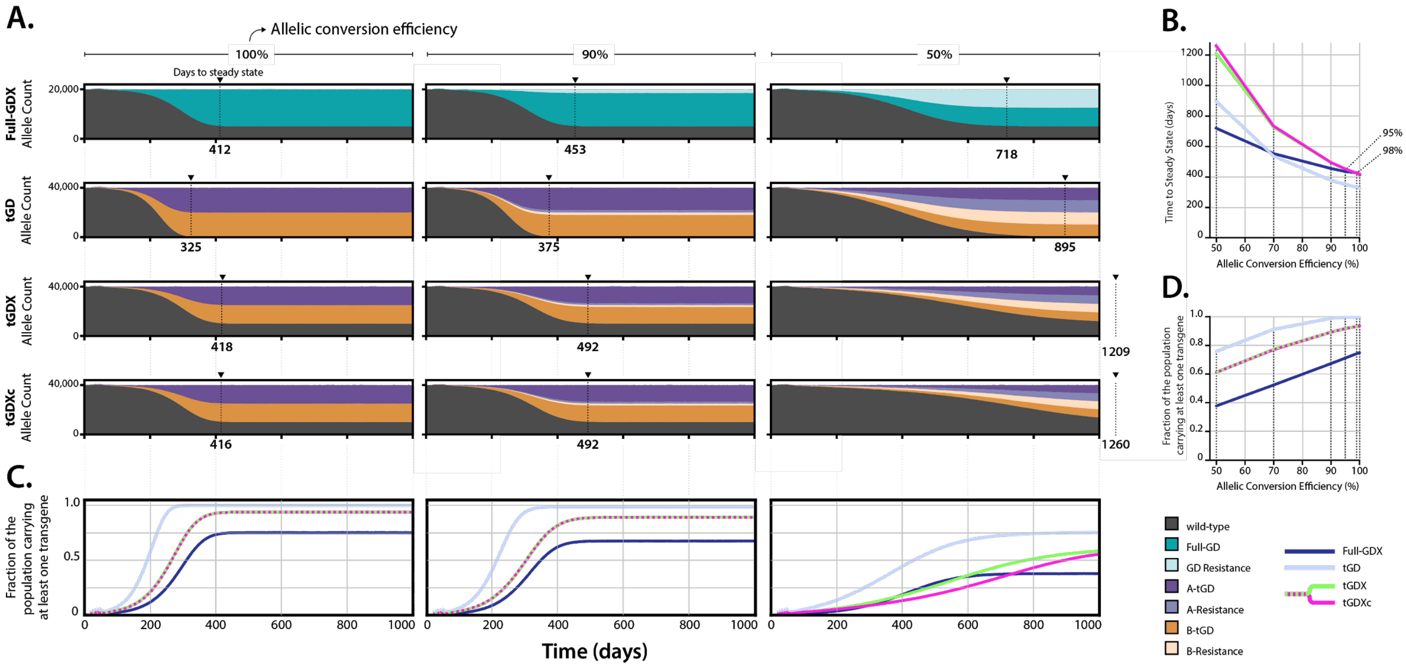


**Supplementary Figure 8 - Mathematical modelling of spread of a variety of trans-complementing and full gene drive systems through *Ae. aegypti* populations.** Model predictions for releases of *Ae. aegypti* mosquitoes homozygous for the trans-complementing system with: i) components linked on an autosome (tGD), ii) components linked on the X chromosome (tGDX), and iii) components unlinked at two loci on the X chromosome (tGDXc). Also modeled are releases of *Ae. aegypti* mosquitoes homozygous for a full gene drive iv) at an X chromosome locus (Full-GDX). Drive systems are parameterized with ballpark parameter estimates for model exploration: i) a cleavage frequency of 100% in females and males, ii) an allelic conversion efficiency given cleavage of 50-100% in females and males, and iii) no fitness costs associated with the Cas9 or gRNA alleles. All resistant alleles are assumed to be in-frame/cost-free. Five weekly releases are simulated consisting of 100 adult males homozygous for each system into a population having an equilibrium size of 10,000 adults. Model predictions were computed using 50 realizations of the stochastic implementation of the MGDrivE simulation framework[2]. **(A)** Stacked allele counts over time are depicted for the Full-GDX, tGD, tGDX, and tGDXc systems for allelic conversion efficiencies of 100%, 90% and 50%. **(B)** Allelic conversion efficiency plotted against time to steady state for the Full-GDX (dark blue), tGD (light blue), tGDX (green), and tGDXc (pink) systems. Autosomal systems spread faster than X-linked systems due to their ability to drive in both sexes. At high allelic conversion efficiencies (90-100%), autosomal systems spread at similar speeds, as do X-linked systems; however as the allelic conversion efficiency declines (50-90%), the tGD and tGDc systems are slowed down to a greater extent than the Full-GD system (compare to **Fig. 6**). Similarly, the tGDX and tGDXc systems are slowed down to a greater extent than the Full-GDX system. (**C**) Fraction of the population carrying at least one transgene over time for the Full-GDX (dark blue), tGD (light blue), tGDX (green), and tGDXc (pink) systems for allelic conversion efficiencies of 100%, 90% and 50%. **(D)** Allelic conversion efficiency plotted against fraction of the population carrying at least one transgene for the Full-GDX (dark blue), tGD (light blue), tGDX (green), and tGDXc (pink) systems. For autosomal systems, while resistant alleles accumulate to similar overall proportions for the tGD, tGDc and Full-GD systems, the tGD and tGDc systems are spread across two loci, and so a higher proportion of individuals have at least one copy of a transgene at equilibrium (for allelic conversion efficiencies <100%). Similarly for X-linked systems, the tGDX and tGDXc systems are spread across two loci, and so a higher proportion of individuals have at least one copy of a transgene, as compared to the Full-GDX system (compare to **Fig. 6**).

**Supplementary Data 1 - Phenotypic analysis of the F2 progeny of tGD(y,e) crosses performed.**

Raw counting data of the F2 progeny phenotypic scoring indicating females and males recovered. Red marker (**DsRed+**), green marker (**GFP+**), both fluorophores (**both**) or no fluorescence (**none**) were scored in order to track Cas9 (red marker) and gRNAs (green marker) transgenes. Transgene inheritance rates in the F2 progeny for each specific tube (marked as **cross#** in the table) were calculated by combining data from males and females. Average inheritance for both markers and the standard deviation are calculated as well. Data is divided in the following file tabs:

1. **Fig. 1C - tGD(y,e) - Raw data:** tGD targeting *yellow* (Cas9-Red) and *ebony* (gRNA-Green).

**Supplementary Data 2 - Phenotypic analysis of the F2 progeny of tGD(y,w) crosses performed.**

Raw counting data of the F2 progeny phenotypic scoring indicating females and males recovered. Red marker (**DsRed+**), green marker (**GFP+**), both fluorophores (**both**) or no fluorescence (**none**) were scored in order to track Cas9 (red marker) and gRNAs (green marker) transgenes. Transgene inheritance rates in the F2 progeny for each specific tube (marked as **cross#** in the table) were calculated by combining data from males and females. Average inheritance for both markers and the standard deviation are calculated as well. The eye color under regular brightfield conditions of each single fly was also scored to distinguish red eye (**w+**), white eye (**w-**) or mosaic eye (**w+/w-**) in both females and males. Data is divided in the following file tabs:

1. **Fig. 2E(A) - tGD(y,w) - Raw data:** tGD targeting *yellow* (Cas9-Red) and *white* (gRNA-Green). Inheritance rates using *vasa* promoter with Cas9 inherited from F0 males.
2. **Fig. 2E(B) - tGD(y,w) - Raw data:** tGD targeting *yellow* (Cas9-Red) and *white* (gRNA-Green). Inheritance rates using *vasa* promoter with Cas9 inherited from F0 females.
3. **Fig. 2E(C) - tGD(y,w) - Raw data:** tGD targeting *yellow* (Cas9-Red) and *white* (gRNA-Green). Inheritance rates using *vasa* promoter with Cas9 and gRNA transgenes inherited from F0 males.
4. **Fig. 2E(D) - tGD(y,w) - Raw data:** tGD targeting *yellow* (Cas9-Red) and *white* (gRNA-Green). Inheritance rates using *vasa* promoter with Cas9 and gRNA transgenes inherited from F0 females.
5. **Supp. Fig. 2 - tGD(w,y) - Raw data:** tGD "swapped" version targeting *yellow* (gRNAs-Red) and *white* (Cas9-Green). Inheritance rates using *vasa* promoter with Cas9 inherited from F0 males.
6. **Supp. Fig. 3(A) - tGD(y,w) - Raw data:** tGD targeting *yellow* (Cas9-Red) and *white* (gRNA-Green). Inheritance rates using *nanos* promoter with Cas9 inherited from F0 males.
7. **Supp. Fig. 3(B) - tGD(y,w) - Raw data:** tGD targeting *yellow* (Cas9-Red) and *white* (gRNA-Green). Inheritance rates using *nanos* promoter with Cas9 inherited from F0 females.

8. **Supp. Fig. 3(C) - tGD(y,w) - Raw data:** tGD targeting *yellow* (Cas9-Red) and *white* (gRNA-Green). Inheritance rates using *nanos* promoter with Cas9 and gRNA transgenes inherited from F0 males.
9. **Supp. Fig. 3(D) - tGD(y,w) - Raw data:** tGD targeting *yellow* (Cas9-Red) and *white* (gRNA-Green). Inheritance rates using *nanos* promoter with Cas9 and gRNA transgenes inherited from F0 females.

**Supplementary Data 3 - Phenotypic analysis of the F2 progeny of drug-inducible tGD(y,w) crosses performed.**

Raw counting data of the F2 progeny phenotypic scoring indicating females and males recovered. Red marker (**DsRed+**), green marker (**GFP+**), both fluorophores (**both**) or no fluorescence (**none**) were scored in order to track Cas9 (red marker) and gRNAs (green marker) transgenes. Transgene inheritance rates in the F2 progeny for each specific tube (marked as **cross#** in the table) were calculated by combining data from males and females. Average inheritance for both markers and the standard deviation are calculated as well. Both spCas9 and DD2-spCas9 were tested in regular food (**TMP-**) and under 80µM Trimethoprim exposure (**TMP+**). Data is divided in the following file tabs:

1. **Supp. Fig. 6B (SpCas9) - tGD(y,w) - Raw data.** tGD targeting *yellow* (Cas9-Red) and *white* (gRNA-Green) using *vasa* promoter and SpCas9.
2. **Supp. Fig. 6B (DD2-SpCas9) - tGD(y,w) - Raw data.** Drug-inducible tGD (DD2-SpCas9) targeting *yellow* (Cas9-Red) and *white* (gRNA-Green) using *vasa* promoter. The eye phenotype of F2 flies were scored to distinguish wild-type, red eye (**w+**), white eye (**w-**) or mosaic eye (**w+/w-**) in both females and males. We also scored flies for yellow body color (**yellow-**) or wild-type body color (**yellow+**).
3. **Supp. Fig. 6C - tGD(y,w) - Raw data.** Daily analysis of gene drive activity in the adult germline using the drug-inducible tGD (DD2-SpCas9) targeting *yellow* (Cas9-Red) and *white* (gRNA-Green) driven by *vasa* promoter. The eye phenotype of F2 flies were scored to distinguish wild-type, red eye (**w+**), white eye (**w-**) or mosaic eye (**w+/w-**) in both females and males. We also scored flies for yellow body color (**yellow-**) or wild-type body color (**yellow+**).
4. **Supp. Fig. 6C - Summary - tGD(y,w) - Raw data.** Summary of the gene drive activation in the adult germline over 10 consecutive days. The eye phenotype of F2 flies were scored to distinguish wild-type, red eye (**w+**), white eye (**w-**) or mosaic eye (**w+/w-**) in both females and males. We also scored flies for yellow body color (**yellow-**) or wild-type body color (**yellow+**).

**Supplementary Data 4 - Phenotypic analysis of the F2 progeny of our gene drive elements with impaired homology arms.**

1. **Fig. 5A (both sides) - tGD(y,w) - Raw data.** tGD targeting *yellow* (Cas9-Red) and *white* (gRNA-Green) with the gRNA transgene lacking 20 nucleotides on both sides.

2. **Fig. 5A (PAM-proximal) - tGD(y,w) - Raw data.** tGD targeting *yellow* (Cas9-Red) and *white* (gRNA-Green) with the gRNA transgene lacking 20 nucleotides in the PAM-proximal side.
3. **Fig. 5A (PAM-distal) - tGD(y,w) - Raw data.** tGD targeting *yellow* (Cas9-Red) and *white* (gRNA-Green) with the gRNA transgene lacking 20 nucleotides in the PAM-distal side.

**References for the Supplementary Information:**


1. Lopez Del Amo, V. *et al.* Small-molecule control of super-Mendelian inheritance in gene drives. doi:10.1101/665620
2. Sánchez C., H. M., Wu, S. L., Bennett, J. B. & Marshall, J. M. MGDrivE: A modular simulation framework for the spread of gene drives through spatially-explicit mosquito populations. doi:10.1101/350488
